# Neurons accumulate disease-specific somatic genomic changes across tau pathologic states in Alzheimer’s disease

**DOI:** 10.1101/2025.05.26.656152

**Authors:** Bowen Jin, Katherine S.-M. Brown, Denis S. Smirnov, Samuel M. Naik, Samantha L. Kirkham, Elizabeth L. Hennessey, Guanlan Dong, Shulin Mao, Samadhi P. Wijethunga, Theresa Connors Stewart, Dennis J. Selkoe, Matthew P. Frosch, Derek H. Oakley, Bradley T. Hyman, August Yue Huang, Michael B. Miller

**Affiliations:** Division of Neuropathology, Department of Pathology, Brigham and Women’s Hospital, Harvard Medical School; Boston, MA, USA; Division of Genetics and Genomics, Department of Pediatrics, Boston Children’s Hospital, Harvard Medical School; Boston, MA, USA; Department of Neurology, Massachusetts General Hospital, Harvard Medical School; Boston, MA, USA; Department of Neurology, Brigham and Women’s Hospital, Harvard Medical School; Boston, MA, USA; Department of Pathology, Massachusetts General Hospital, Harvard Medical School; Boston, MA, USA; Broad Institute of MIT and Harvard; Cambridge, MA, USA

**Author notes:** These authors contributed equally to this work (Bowen Jin, Katherine S.-M. Brown). These authors jointly supervised this work (August Yue Huang, Michael B. Miller).

## Abstract

Tau deposition within neurons marks Alzheimer’s disease (AD) neuropathology, suggesting that tau may contribute to cellular dysfunction and death. In AD, somatic mutations accumulate in neurons, with features that suggest deleterious effects on cellular function. To examine the relationship between tau and somatic mutation, we isolated neurons according to tau pathology and performed single-cell whole-genome sequencing on tau+, tau–, and tau-agnostic neurons from 13 individuals with AD, as well as neurons from 15 control individuals. We found that AD neurons, regardless of tau status, exhibited an increased burden of somatic single-nucleotide variants (sSNVs) and insertions and deletions (sIndels). Mutational signature analyses revealed disease-specific patterns, including sIndels characterized by two-basepair (2bp) deletions, indicating shared mutagenic mechanisms in AD neurons across tau pathologic cell states. Somatic mutations are associated with tissue-wide tau pathology, suggesting that tangles do not confer cell-autonomous genotoxicity to neurons and that non-tangle components drive somatic mutation in AD.

## Main Text

Alzheimer’s disease (AD) is a progressive and fatal neurodegenerative disorder characterized by pathologic hallmarks of extracellular amyloid-β (Aβ) plaques and intracellular tau aggregates in neurofibrillary tangles (NFTs). While Aβ plaque accumulation begins decades before symptom onset, tau NFTs are spatiotemporally associated with neurodegeneration and cognitive decline, such that tau Braak staging is the strongest pathologic predictor of AD clinical dementia (*1–4*). Despite the critical role of NFTs in defining AD pathology, the precise mechanisms by which tau contributes to AD pathogenesis remain poorly understood. The correlation of NFTs with neuronal loss potentially implicates tau in neuronal death (*5*), though cell death exceeds neuronal loss and a protective NFT role has also been postulated (*6*, *7*).

Somatic mutations accumulate in healthy human cells, including in neurons during aging (*8–11*). We previously found that AD neurons demonstrate an increased burden of somatic single-nucleotide variants (sSNVs) relative to neurotypical controls, with distinct disease-related mutational signatures partially driven by oxidative damage (*12*). However, the relationship between AD-related protein misfolding pathology and these somatic genomic changes has not been examined.

Despite the strong association with AD progression, NFT pathology varies between brain regions and shows heterogeneity at the single-cell level, burdening only a subset of neurons within affected brain regions. The etiology and pathogenic implications of this heterogeneity have yet to be resolved. Here, we interrogated NFT pathology and somatic genomic changes in single neurons to test whether AD tau pathologic state–as defined by the presence of tau NFTs–is associated with disease-specific somatic mutations. To achieve this, we developed a method to isolate single neuronal nuclei on the basis of NFT pathology and applied strand-agnostic and duplex-strand single-cell whole genome sequencing (scWGS) methods to profile two forms of somatic mutations, sSNVs and somatic insertions and deletions (sIndels), with strand-specific resolution. This approach enabled us to profile somatic genomic variants across tau pathologic states and evaluate for cell-autonomous changes related to tau pathology.

## Results

### Somatic mutations in AD neurons correlate with tissue-wide tau pathology

We previously found that neurons accumulate sSNVs with age, and that sSNV accumulation is elevated in AD neurons (*12*). To examine multiple classes of somatic genomic alterations, we reanalyzed our scWGS data on control and AD neurons isolated from human brain prefrontal cortex (PFC) using a new tool that enables the analysis of both sSNVs and sIndels (*8*).

Consistent with our previous study, we found that sSNVs are elevated in AD neurons relative to controls (by 228 sSNVs per neuron, *P* = 5.72×10^-6^, two-tailed Wilcoxon test, Fig. 1A). Further, while sIndels accumulate minimally in controls, AD neurons show a pronounced increase (mean = 618 sIndels per neuron, median = 107, *P* = 4.83×10^-4^, two-tailed Wilcoxon test), with some AD neurons harboring up to 8,000 sIndels (Fig. 1A).

**Fig. 1.**
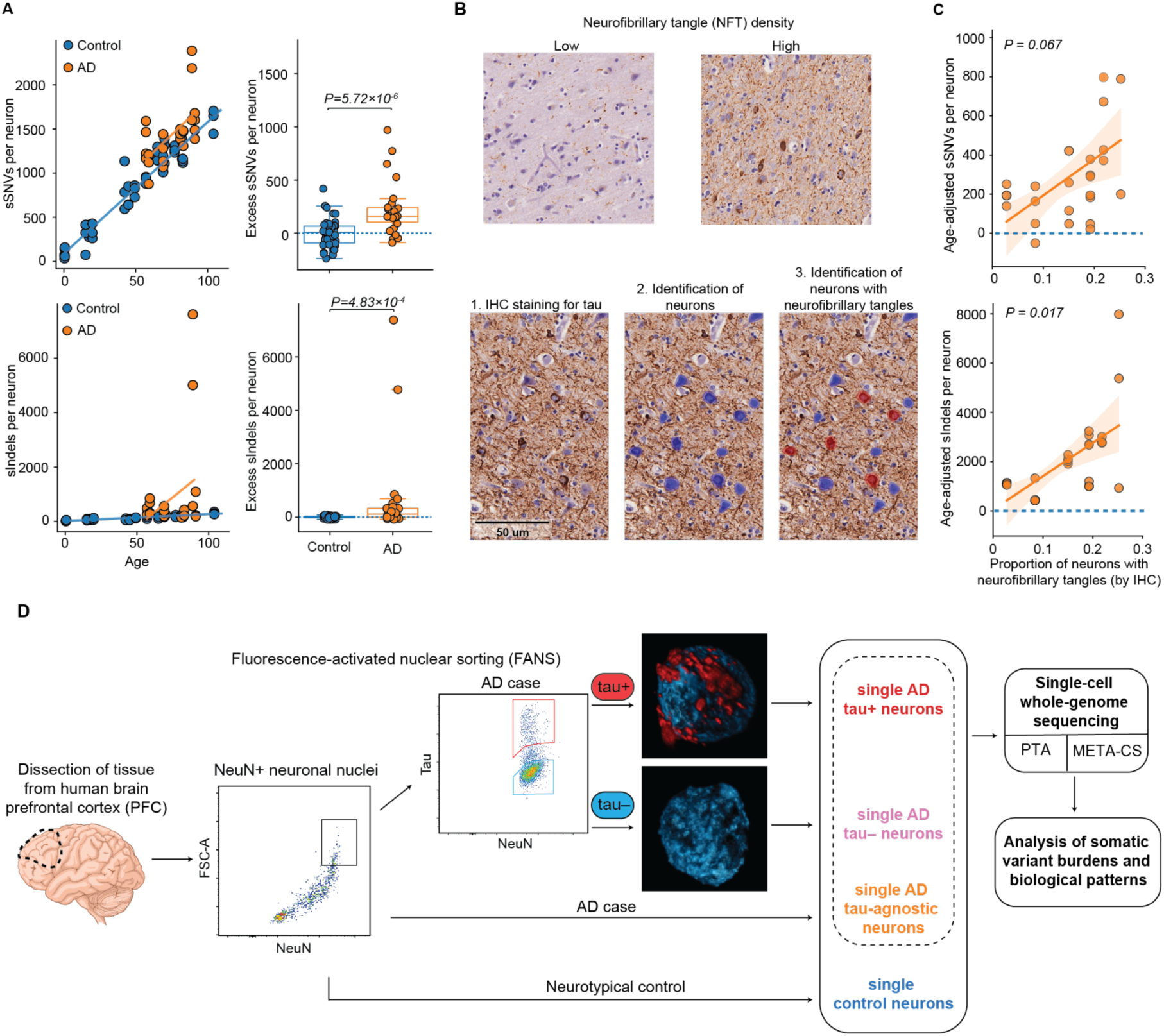
Somatic mutations in AD neurons relate to tau pathology. **(A)** Increased sSNV and sIndel burden in AD neurons compared to neurotypical controls. sSNV burden (from each neuron as a point) analyzed by SCAN2 is shown against age. AD neurons show a significant excess of sSNV (*P* = 5.72×10^-6^, two-tailed Wilcoxon test) and sIndel (*P* = 4.83×10^-4^, two-tailed Wilcoxon test) burdens compared to neurotypical controls. **(B)** Quantitative immunohistochemistry (IHC) of neurofibrillary tangles (NFTs) in neurons in AD brain, using QuPath to identify and determine the proportion of NFT-bearing neurons. **(C)** Relationship between tissue proportion of NFT-bearing neurons and neuronal somatic mutation burden. Age-adjusted sSNV and sIndel burdens are fitted against the proportion of neurons with NFTs, as identified by IHC, using a LME model (*P* = 0.067 and *P* = 0.017, respectively). The burden of somatic mutations per neuron correlates with tissue-wide NFT pathology. **(D)** Study design and single-cell whole-genome sequencing experimental approach. Nuclei are isolated from postmortem human brain prefrontal cortex tissue and undergo fluorescence-activated nuclear sorting (FANS) for the neuronal nuclear marker NeuN to obtain a neuronal population. Such tau-agnostic neurons are further sorted by their tau pathologic state using FANS. The three populations of AD neurons—tau+, tau–, and tau-agnostic–were co-analyzed for each individual. All four populations of neurons were profiled with two single-cell whole genome sequencing (scWGS) methods, primary template-directed amplification (PTA), and multiplexed end-tagging amplification of complementary strands (META-CS). Amplified genomic DNA then underwent whole-genome sequencing to identify somatic mutations. Somatic genomic changes of tau-defined neurons were compared to tau-agnostic AD and neurotypical control neurons.

We then sought to examine potential mechanisms that may contribute to somatic mutation accumulation in AD. First, we analyzed neuropathologic features for correlations with mutation burden. NFTs form the basis of the pathologic staging scheme that most strongly predicts clinical dementia, yet they burden only a subset of neurons in affected brain tissue. Also, even in advanced Braak stages, the proportion of NFT-bearing neurons within a given brain region can vary considerably (*13*). The etiology and pathogenic implications of this heterogeneity are unresolved.

To investigate the dependence of sSNVs and sIndels with tissue-wide pathology, we performed immunohistochemistry (IHC) to label NFTs, illustrating a range of densities in our AD cohort. We quantified the proportion of neurons bearing NFTs across the PFC of each case (Fig. 1B), and assessed their relationship with somatic mutation burdens. We found that sIndel burdens from single neurons correlate positively with the tissue-wide proportion of NFT-bearing neurons (*P* = 0.017, linear mixed-effects (LME) model, Fig. 1C), while sSNVs showed a non-significant positive relationship (*P* = 0.067; Fig. 1C). Semi-quantitative assessments of amyloid-β plaques and NFTs, retrieved from pathology reports, also showed trends toward greater mutational burden, albeit non-significant (fig. S1). These observations suggest that tau NFTs may influence somatic mutation accumulation in AD.

### Isolation of neuronal nuclei by tau pathologic state for single-cell genome sequencing

After observing that increased somatic mutations in AD aligned with tissue-level tau NFT abundance (Fig. 1C), we turned to examine the influence of tau at the single-neuron level. We sought to compare the somatic genomic landscape of AD neurons with distinct NFT pathologic states to provide insight into the mechanisms by which tau aggregation may contribute to AD pathogenesis. To do so, we developed a fluorescence-activated nuclear sorting (FANS) method to separate the nuclei of neurons by their NFT pathologic state, enabling us to compare the somatic genomes between NFT-bearing and NFT-free neurons (Fig. 1D).

To profile the NFT pathologic state, we extracted nuclei from postmortem human brain tissue and isolated neurons with NeuN. We then distinguished nuclei of NFT-bearing neurons (tau+) from NFT-free neurons (tau–) with anti-tau antibodies. We compared these tau-sorted neurons with AD “tau-agnostic” neurons not profiled for tau and with non-diseased neurons from neurotypical control individuals. We performed scWGS on neurons using two distinct genome amplification methods: primary template-directed amplification (PTA) (*14*) and multiplexed end-tagging amplification of complementary strands (META-CS) (*10*). We utilized PTA as a well-established method to assess the burden of somatic mutations (*8*). We compared tau-agnostic, tau+, and tau– neurons from the same individuals with AD as internal comparisons to minimize confounding factors, and we compared AD neurons with controls to distinguish somatic genomic changes driven by age from AD.

### Somatic SNVs accumulate in AD neurons independently of tau NFT pathology

Based on our observation of increased somatic mutations in AD and their positive association with tissue-wide NFT pathology (Fig. 1A, C), we hypothesized that NFT-bearing AD neurons would have a greater somatic mutation burden than AD neurons lacking tau NFT pathology. We performed PTA-based scWGS on 51 tau+ neurons and 26 tau– neurons from 7 AD cases and analyzed them alongside previously generated data from 28 AD tau-agnostic neurons and 49 neurotypical control neurons from individuals ranging from 0.7 to 104 years old (Table 1).

**Table 1.** Case information with number of neurons analyzed using two scWGS methods.

| Case ID | Age (years) | Sex | Diagnosis | Neurons, PTA-amplified |  |  | Neurons, META-CS-amplified |  |  |
| --- | --- | --- | --- | --- | --- | --- | --- | --- | --- |
| Controls |  |  |  |  |  |  |  |  |  |
| Younger neurotypical controls |  |  |  |  |  |  |  |  |  |
| 1278 | 0.4 | M | Normal | 3 |  |  | 10 |  |  |
| 5817 | 0.6 | M | Normal | 3 |  |  | 17 |  |  |
| 4638 | 15.1 | F | Normal | 3 |  |  | 9 |  |  |
| 1465 | 17.5 | M | Normal | 4 |  |  | 10 |  |  |
| 5559 | 19.8 | F | Normal | 3 |  |  | - |  |  |
| 4643 | 42.2 | F | Normal | 3 |  |  | - |  |  |
| 5087 | 44 | M | Normal | 3 |  |  | 9 |  |  |
| 936 | 49.2 | F | Normal | 3 |  |  | 10 |  |  |
|  |  |  |  | 25 |  |  | 65 |  |  |
| Aged neurotypical controls |  |  |  |  |  |  |  |  |  |
| 5451 | 57 | F | Normal | 3 |  |  | 17 |  |  |
| 5666 | 65 | M | Normal | 3 |  |  | 20 |  |  |
| 5943 | 69 | M | Normal | 3 |  |  | 10 |  |  |
| 5572 | 70 | F | Normal | 3 |  |  | 10 |  |  |
| 5219 | 77 | F | Normal | 3 |  |  | 10 |  |  |
| 5657 | 82.2 | M | Normal | 3 |  |  | 9 |  |  |
| 5823 | 82.7 | F | Normal | 3 |  |  | 19 |  |  |
| 4976 | 104 | F | Normal | 3 |  |  | 18 |  |  |
|  |  |  |  | 24 |  |  | 113 |  |  |
| Alzheimer's disease |  |  |  |  |  |  |  |  |  |
|  |  |  |  | Tau-agnostic | Tau+ | Tau– | Tau-agnostic | Tau+ | Tau– |
| 1353 | 57 | F | AD (Braak VI) | 4 | 6 | 4 | - | 10 | 10 |
| 1647 | 59 | F | AD (Braak VI) | 4 | 9 | 4 | 10 | 8 | 10 |
| 2210 | 66 | F | AD (Braak VI) | - | - | - | 10 | - | - |
| 2208 | 69 | F | AD (Braak VI) | 4 | 8 | 4 | 10 | 19 | 10 |
| 2203 | 75 | F | AD (Braak VI) | - | - | - | 10 | 9 | 10 |
| 2205 | 79 | F | AD (Braak VI) | - | - | - | 8 | - | - |
| 1456 | 81 | M | AD (Braak VI) | 5 | 8 | 4 | 9 | 10 | 10 |
| 2219 | 82 | M | AD (Braak VI) | - | - | - | 9 | 10 | 18 |
| 2207 | 83 | M | AD (Braak VI) | 3 | 4 | 4 | 9 | - | - |
| 1995 | 89 | F | AD (Braak V) | 4 | 8 | 3 | 9 | 9 | 10 |
| 2204 | 89 | F | AD (Braak VI) | - | - | - | 10 | - | - |
| 2211 | 90 | M | AD (Braak VI) | - | - | - | 10 | 10 | - |
| 1828 | 91 | F | AD (Braak VI) | 4 | 8 | 3 | - | 8 | 18 |
|  |  |  |  | 28 | 51 | 26 | 104 | 93 | 96 |

Surprisingly, we found no significant difference in the total sSNV burden between tau+ and tau– neurons (*P* = 0.75, LME model, Fig. 2A). We further calculated the excess sSNV burden for each pathologic state, adjusting for age using linear regression. Tau+ neurons showed a marginally lower median sSNV burden than tau– neurons, which did not reach statistical significance (*P* = 0.313, two-tailed Wilcoxon test, Fig. 2B), which could suggest that tau tangles exert a slight protective effect against mutations. However, all AD neurons, independently of tau status, exhibited an excess in sSNVs relative to controls (tau+ vs. controls: *P* = 5.73×10^-3^; tau– vs. controls: *P* = 1.96×10^-3^, two-tailed Wilcoxon test, Fig. 2B). We utilized three different antibodies to identify tau+ pathologic state that yielded comparable findings (fig. S2), and thus we combined their data in our analyses. Of note, we profiled neurons isolated using total tau as well as phospho-specific antibodies, such that the targeted tau aggregates represent NFT state broadly, rather than hyperphosphorylation alone.

**Fig. 2.**
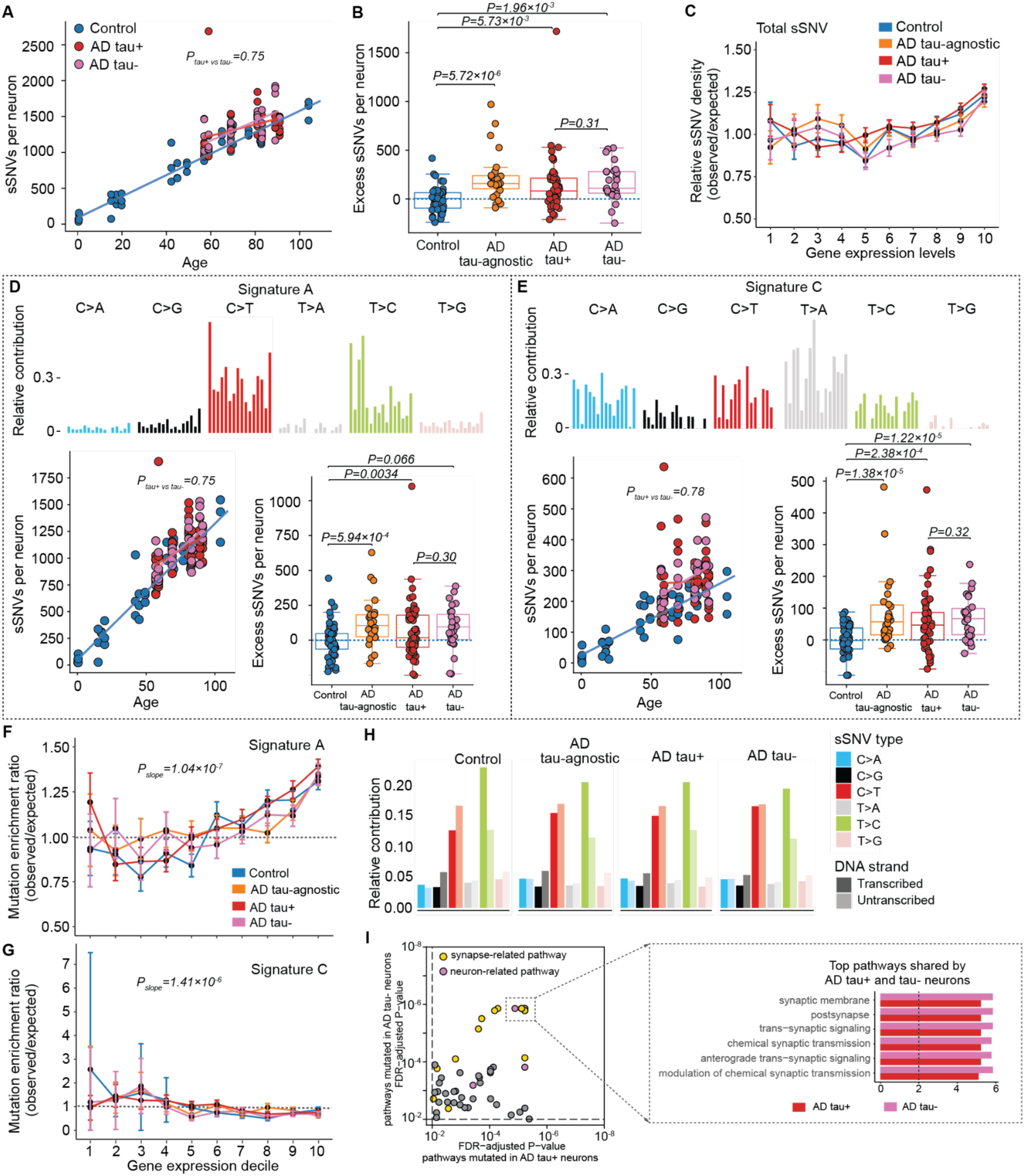
Somatic SNV mutational burden and patterns in single AD neurons across tau pathologic states. **(A)** sSNV burden as a function of age in controls and AD neurons with distinct tau states. sSNVs of tau+ AD, tau– AD, and control neurons are fitted against age using a LME model (tau+ vs controls: *P* = 2.74×10^-3^; tau– vs controls: *P* = 2.9×10^-3^; tau+ vs tau–: *P* = 0.75). AD tau+ and tau– do not significantly differ in mutational burden, although sSNVs are increased for all AD neurons, regardless of tau pathologic state, relative to controls. **(B)** Excess sSNVs per neuron, calculated as the residual value from linear regression adjusting for age. Tau+ and tau– neurons from AD show similar increases in sSNVs compared to age-matched control neurons (tau+ vs. controls: *P* = 5.73×10^-3^; tau– vs. controls: *P* = 1.96×10^-3^, two-tailed Wilcoxon test). Tau+ and tau– neurons show no significant difference in excess sSNV burden (*P* = 0.313). (**C**) Genomic sSNV density as a function of gene expression. Overall, SNV density increases in genes with higher expression levels. Genes are grouped into deciles by their expression in neurons. Data points represent the ratio of observed to expected sSNV density. Error bars indicate 95% confidence intervals. **(D, E)** sSNV mutational signature analysis, showing contributions of neuronal Signatures A (D) and C (E) to sSNV burden by age and as excess versus control neurons. (Signature A: tau+ vs. controls: *P* = 0.0034; tau– vs. controls: *P* = 0.066, two-tailed Wilcoxon test). Tau+ vs. tau– shows no significant difference in excess Signatures A burden (*P* = 0.3). Signature C: tau+ vs. controls: *P* = 2.38× 10^-4^; tau– vs. controls: *P* = 1.22×10^-5^, two-tailed Wilcoxon test). Tau+ vs. tau– shows no significant difference in excess Signature C burden (*P* = 0.32). **(F, G)** sSNV density by signature as a function of gene expression, for Signatures A (F) and C (G). Genes are grouped into deciles by their expression in neurons. Error bars indicate the 95% confidence interval. **(H)** sSNVs by DNA strand template status across clinical conditions and tau pathologic states. Relative contributions of the transcribed and untranscribed strands are shown for each nucleotide change. **(I)** Ontology analysis of genes mutated in AD tau+ and AD tau– neurons, with pathways enriched in each population (FDR < 0.01). Synaptic and neuronal-related pathways are highlighted in yellow and purple. The rectangle denotes the top pathways (6/10) shared by both neuronal populations.

To determine whether processes driving sSNV accumulation in AD neurons differ by tau state, we first examined the relationship between transcription and somatic mutation, and observed that sSNVs occur more frequently in highly expressed genes, independently of disease status and tau pathologic state (*P* = 4.49×10^-4^, linear model, Fig. 2C). We next performed mutational signature analysis. First, we decomposed sSNVs into the previously reported neuronal Signatures A and C (*9*) (Fig. 2D, E). Signature A closely resembles signature SBS5 from the COSMIC cancer database, a clock-like signature associated with genome aging. While the contribution of Signature A mutations does not differ by tau state, Signature A shows a modest increase in tau+, tau–, and tau-agnostic AD neurons relative to age-matched controls (tau+ vs. controls: *P* = 0.0034; tau– vs. controls: *P* = 0.066, tau-agnostic vs. controls: *P* = 5.94×10^-4^; two-tailed Wilcoxon test, Fig. 2D). Moreover, we observe similar influences of transcriptional activity on Signature A in control neurons and AD neurons, with the burden of sSNVs attributed to Signature A positively correlated with gene expression (Fig. 2F).

AD neurons of both tau states demonstrate a substantial increase in Signature C compared to controls, largely accounting for the excess sSNVs observed in AD (tau+ vs. controls: *P* = 2.38×10^-4^; tau– vs. controls: *P* = 1.22×10^-5^, two-tailed Wilcoxon test, Fig. 2E). Signature C is characterized by C>A substitutions, which are associated with oxidative damage to guanine nucleotides, and has also been linked to transcription-coupled nucleotide excision repair (*12*). Notably, Signature C burdens do not differ between tau+ and tau– AD neurons (*P* = 0.32, LME model), suggesting that AD neurons with differential NFT pathology are indiscriminately affected by oxidative stress in AD pathogenesis. In contrast to Signature A, we observe a negative correlation between Signature C mutations and gene expression (*P* = 1.41×10^-7^, linear model, Fig. 2G), supporting their association with transcription-coupled nucleotide excision repair which preferentially repairs mutations in highly expressed genes.

Because transcriptional influences on somatic mutations can produce asymmetry on paired DNA strands, we distinguished sSNV sites by template status, and observed no difference between tau+ and tau– neurons (Fig. 2H). To further examine the relationship with gene expression, we performed gene ontology analysis to identify pathways with genes bearing mutations, finding that genes for neuronal and synaptic function are significantly enriched in both tau+ and tau– neurons, again pointing to common somatic mutational features regardless of tau pathologic state in AD (Fig. 2I).

To further investigate the strand-specificity of sSNVs in AD, we employed the duplex-strand scWGS method META-CS (*10*). We performed META-CS on 110 tau+ AD neurons, 110 tau– AD neurons, 110 tau-agnostic AD neurons, and 80 control neurons, with paired PTA scWGS data. We classified single-nucleotide variants as either double-stranded mutations (dsSNVs) or single-stranded DNA damage lesions (ssSNVs), depending on whether the alteration was supported by reads from both the Watson and Crick strands or from only one of the two strands, and then generated mutational signatures for each category (fig. S3A). Decomposition of PTA sSNV data using these dsSNV and ssSNV signatures showed a mean 91.0% double-stranded contribution (fig. S3B). These findings indicate that PTA-derived sSNV calls are predominately double-stranded, which is corroborated by the similar neuron aging sSNV rates observed between duplex-strand approaches (*10*, *11*) and PTA (*8*). This strand-aware analysis supports the sSNV burden and signature analyses that indicate that sSNVs accumulate similarly across tau pathologic states in AD.

### Somatic deletions accumulate in AD neurons across tau pathologic states

As with sSNVs, our observation that increased sIndels in AD are associated with tissue-level tau pathology (Fig. 1A, C) prompted us to examine sIndels in single neurons with distinct tau NFT pathologic states. Analyzing PTA scWGS data, we observed that the dramatic increase of thousands of excess sIndels in AD neurons compared to normal aging occurs in each tau pathologic state group (Fig 3A, B; tau+ vs. controls: *P* = 2.9×10^-4^; tau– vs. controls: *P* = 1.05×10^-5^; tau-agnostic vs. controls: *P* = 4.83×10^-4^, two-tailed Wilcoxon test). Notably, the sIndel burden does not differ significantly between AD neurons of different tau states, although tau+ neurons exhibit a marginally lower sIndel burden relative to tau– neurons. (*P* = 0.19, two-tailed Wilcoxon test, Fig. 3B). We next investigated how sIndels may be impacted by gene transcription, and observed a positive correlation with gene expression in neurons isolated from neurotypical control individuals (*P* = 4.49×10^-4^, linear model, Fig. 3C), consistent with the relationship observed for sSNVs. However, transcriptional activity has a more modest influence on sIndel density in AD neurons, a relationship that is unaltered by the presence of tau NFTs. Like with sSNVs, AD-related sIndels are less positively correlated with transcription than aging-related variants. These findings suggest that sIndels may arise through different mechanisms in control versus AD neurons.

**Fig. 3.**
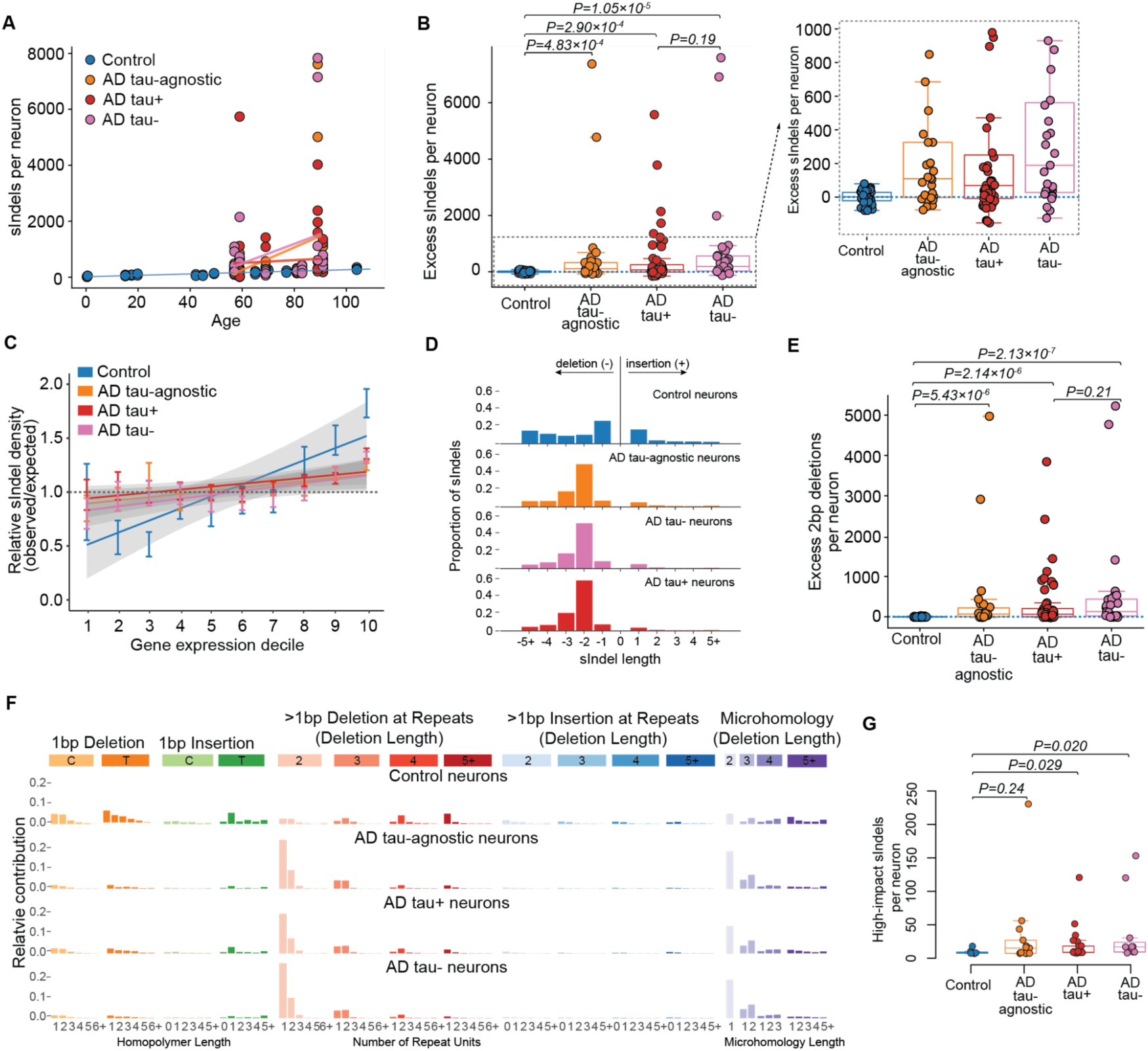
Somatic Indel burden and spectra in single AD neurons across tau pathologic states. **(A)** sIndels as a function of age in control and AD neurons with distinct tau states. **(B)** Excess sIndels per neuron, calculated as the residual value from linear regression adjusting for age. AD contributes a significant excess of sIndels relative to control neurons independently of tau state (tau+ vs. controls: *P* = 2.9×10^-4^; tau– vs. controls: *P* = 1.05×10^-5^, tau+ vs. tau–: *P* = 0.19, two-tailed Wilcoxon test). **(C)** sIndel density as a function of gene expression. sIndel density in control neurons increases in genes with higher expression levels (*P* = 7.94×10^-4^, linear model). Error bars indicate 95% confidence intervals. **(D)** Contribution of sIndels by length in control and AD neurons across tau states. Negative length is used for deletions, and positive length is used for insertions. **(E)** Excess 2bp deletion per neuron, calculated as the residual value from linear regression adjusting for age. AD neurons contribute an excess of 2bp deletions relative to controls independently of tau state (tau+ vs controls: *P* = 2.14×10^-6^; tau– vs controls: *P* = 2.13×10^-7^, two-tailed Wilcoxon test). Tau+ vs tau– show no significant difference in the excess 2bp deletion burden (*P* = 0.21). **(F)** sIndel mutational spectra of control and AD neurons across tau pathologic states. Spectra highlights somatic deletion contexts common to AD neurons across tau states. **(G)** AD neurons harbor more high-impact sIndels, defined as sIndels introducing frameshift, stop gain, and stop loss, than age-matched controls.

Accordingly, we sought to gain further insight into the processes driving sIndel accumulation by examining the lengths of sIndels present in control and AD neurons. We observed that sIndels in AD neurons are predominantly 2bp and 3bp deletions, in contrast to control sIndels where single base deletions and insertions are most abundant (Fig. 3D). 2bp deletions account for over 40% of the total sIndel burden in AD, a pronounced increase relative to the 20% contribution observed in controls. Both tau+ and tau– neurons show a significantly higher total 2bp deletion burden compared to control neurons (tau+ vs. controls: *P* = 2.14×10^-6^; tau– vs. controls: *P* = 2.13×10^-3^, two-tailed Wilcoxon test, Fig. 3E). As our findings suggest that 2bp deletions are the primary drivers of excess sIndel accrual in AD, we examined the mutational spectra of these sIndels for further insights. We observed that the 2bp-deletion motif shown by AD neurons predominantly occurs in the context of 1 or 2 repeat units and in regions of microhomology (Fig. 3F). Notably, sIndel mutational burden and spectra do not show substantial differences by tau NFT pathology, indicating that the processes driving sIndel accrual in AD are independent of tau aggregation (*P* = 0.21, two-tailed Wilcoxon test, Fig. 3E).

To assess the potential deleterious effects on cellular function from increased sIndel accumulation in AD, we examined the abundance of high-impact indels, defined as sIndels introducing frameshifts, stop codon gains, and stop codon losses, and found a substantial increase in AD neurons (Fig. 3G and fig. S4). The effects of high-impact sIndels on genes associated with neuronal function represent one mechanism by which sIndels may contribute to AD pathogenesis. Interestingly, the excess of high-impact sIndels in AD neurons does not vary by tau state, again suggesting that this pathogenic effect is not directed by tau aggregation.

### Novel signatures of deletion variants in AD neurons

Since our findings suggest that increases in high-impact sIndels may contribute to AD pathogenesis, we performed signature analyses to investigate the processes underlying disease-related sIndel accumulation. We began with *de novo* sIndel mutational signature analysis and identified two signatures: one associated with aging (ID-Age) and the other specifically enriched in AD (ID-AD) (Fig. 4A and fig. S5). To understand the underlying mechanistic elements in these new signatures, we examined them for features from previously characterized sIndel patterns from cancer datasets. We first performed decomposition using COSMIC sIndel signatures (*15*). While we identified several COSMIC signatures contributing to the aging-associated signature ID-Age, no COSMIC signature strongly fit the AD-associated ID-AD (fig. S5B).

**Fig. 4.**
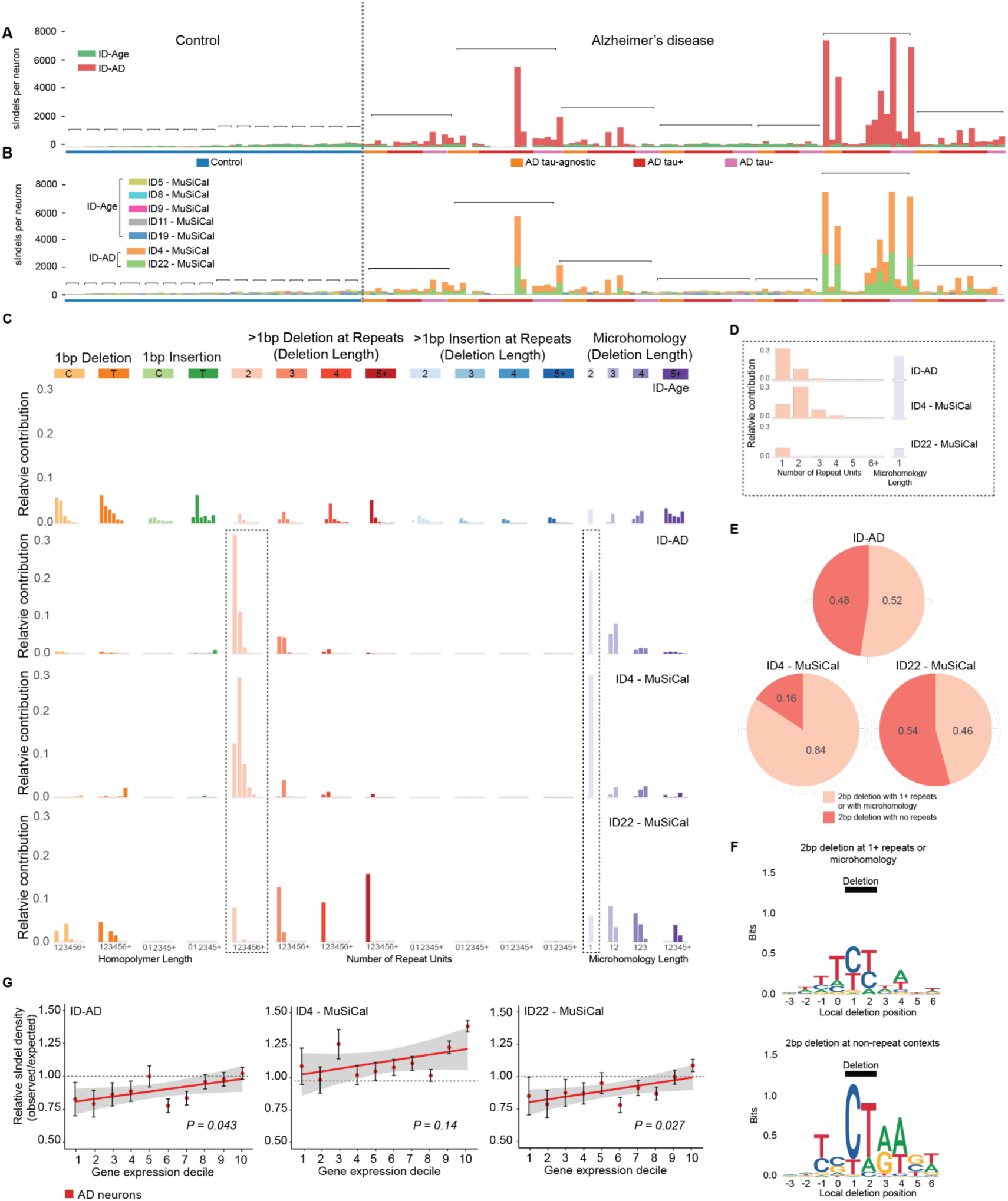
Somatic Indels show deletion mutational signatures in AD neurons across tau pathologic states. Single-neuron sIndel mutational spectra were decomposed *de novo* into two signatures, ID-AD and ID-Age, which underwent further mutational signature analysis to identify potential processes contributing to AD pathogenesis. **(A)** Decomposition of single-neuron PTA data with *de novo* signatures ID-AD and ID-Age. Control neurons show primarily ID-Age, while ID-AD is the primary contributor of excess sIndels in AD tau-agnostic, tau+, and tau– neurons. Brackets identify neurons from each individual, with colors denoting the tau pathologic state for AD neurons. **(B)** ID-AD can be resolved into MuSiCal signatures ID4 and ID22. **(C)** Indel mutational spectra of signatures ID-Age, ID-AD, MuSiCal ID4, and MuSiCal ID22. **(D)** Primary 2bp deletion spectra for ID-AD, MuSiCal ID4, and MuSiCal ID22. ID4 deletions are enriched in genomic contexts of 2 or more repeat units and microhomology and ID22 deletions enriched in non-repeat contexts and microhomology. **(E)** Pie charts for the proportions of different types of 2 bp deletions. The frequencies of 2 bp deletions without repeats, with one or more repeats, and with microhomology were calculated separately for ID-AD, ID4 and ID22. **(F)** Motif analysis for different types of 2bp deletions. **(G)** Density of *de novo* signature ID-AD and MuSiCal signatures ID4 and ID22 as a function of gene expression. All three AD populations were combined for this analysis, as sIndel mutational spectra are consistent across tau states.

We next utilized a different set of sIndel signatures, MuSiCal (*16*), which are also derived from cancers and are highly similar to 16 of the 18 COSMIC signatures, while additionally offering 9 distinct signatures for more granular decomposition. Signature analysis decomposed ID-AD into two MuSiCal signatures: ID4, which is nearly identical to COSMIC ID4, and ID22 (Fig. 4B). A combination of MuSiCal signatures ID4 and ID22 reconstructs a profile highly similar to ID-AD (cosine similarity = 0.82), outperforming any individual sIndel signature from MuSiCal or COSMIC alone (cosine similarity = 0.774 for MuSiCal ID4 and 0.708 for COSMIC ID4). These signatures collectively contribute to the excess of 2bp deletions observed in AD, with ID4 deletions enriched in genomic contexts with repeats (i.e., 2 or more repeat units) and microhomology, whereas ID22 deletions are enriched in non-repeat contexts (Fig. 4C-E). ID4 and ID22 both show specificity for AD, with minimal signal in control neurons.

As one potential mechanism, ID4 has been linked to topoisomerase I (TOP1)-mediated ribonucleotide excision repair (RER) (*17*). The concentration of 2bp deletions in repeat regions at TNT motifs (Fig. 3F) is consistent with a TOP1-dependent mechanism, which may link elements of the ID-AD signal with RER-associated deletions. In contrast, ID22 has not been linked with a specific etiology, and its unique deletion contexts suggest mechanisms distinct from ID4. ID22 features non-2bp deletions, including 3bp deletions, that are not prominent in RER-related mutagenesis. Furthermore, ID22 and ID-AD deletions are concentrated at non-repeat sites, in contrast to RER-related deletions (Fig. 4E), suggesting that many AD-related deletion variants may derive from mechanisms distinct from RER-TOP1. Contribution from a non-RER-mediated mechanism is further supported by motif analysis, which reveals that 2bp deletions in non-repeat contexts show broader NT motifs, most commonly CT deletions (Fig. 4F, fig. S8).

Our analysis suggests that the AD somatic deletions observed in the ID-AD signature may arise from both RER-associated and RER-independent mechanisms. Apart from RER, DNA polymerase strand slippage is a common deletion mechanism in dividing cells, but appears to be less relevant for post-mitotic neurons. As another such potential RER-independent mechanism that may be more relevant in neurons, nitrated polycyclic aromatic hydrocarbons (nitro-PAHs) have been reported to cause multi-base deletions (*18*).

Given the potential for RER and other transcription-coupled processes to cause sIndels, we next investigated the correlation between the burden of AD sIndels and gene expression. We found that ID-AD sIndel density increases with gene expression (Fig. 4G, *P* = 0.043, linear model), and then examined the components of ID-AD, MuSiCal signatures ID4 and ID22. We found that ID-AD’s correlation with gene expression is most strongly derived from ID22, while ID4 shows a non-significant association with gene expression (*P* = 0.027 for ID22; *P* = 0.14 for ID4, linear model, Fig. 4G and fig. S6). ID4 and ID22’s differing strengths of association with gene expression, in addition to their unique spectra, indicate that they may arise from distinct mutagenic mechanisms. However, the overall trends of correlation with gene expression suggests that transcription-associated mutagenesis contributes to AD-associated sIndels.

### sIndels in AD neurons are predominantly single-stranded

To investigate the strand-specificity of AD sIndels, we supplemented the PTA strand-agnostic assessment of sIndel variants with duplex-strand META-CS scWGS. Strand-resolved sIndel analysis revealed that control and AD neurons possess distinct double-stranded (dsIndel) and single-stranded sIndel (ssIndel) mutational signatures (Fig. 5A-B). Notably, the AD signatures are enriched in 2bp deletions in the context of 1-2 repeat units and microhomology, fitting the pattern observed from PTA (Fig. 3F). We quantified the contributions of these 2bp deletions, observing significantly increased proportions in ssIndels in each AD pathologic state over controls (Fig. 5C). dsIndels also showed increased 2bp deletion contributions in each tau state.To examine the potential mechanisms underlying ssIndels and dsIndels in AD, we decomposed their signatures into the reference MuSiCal Indel signatures (Fig. 5D). We identified ID4 and ID22 as the primary contributors to ssIndels in AD neurons, with each as a partial contributor to AD dsIndels. ID12, the most substantial contributor to control neuron ssIndels, also occurred in AD ssIndels. Likewise, the dsIndels in AD show substantial similarities to control dsIndels, including significant signals from clock-like signature ID5 characterized by thymidine deletions, tobacco-associated ID3 characterized by cytidine deletions, and others.

**Fig. 5.**
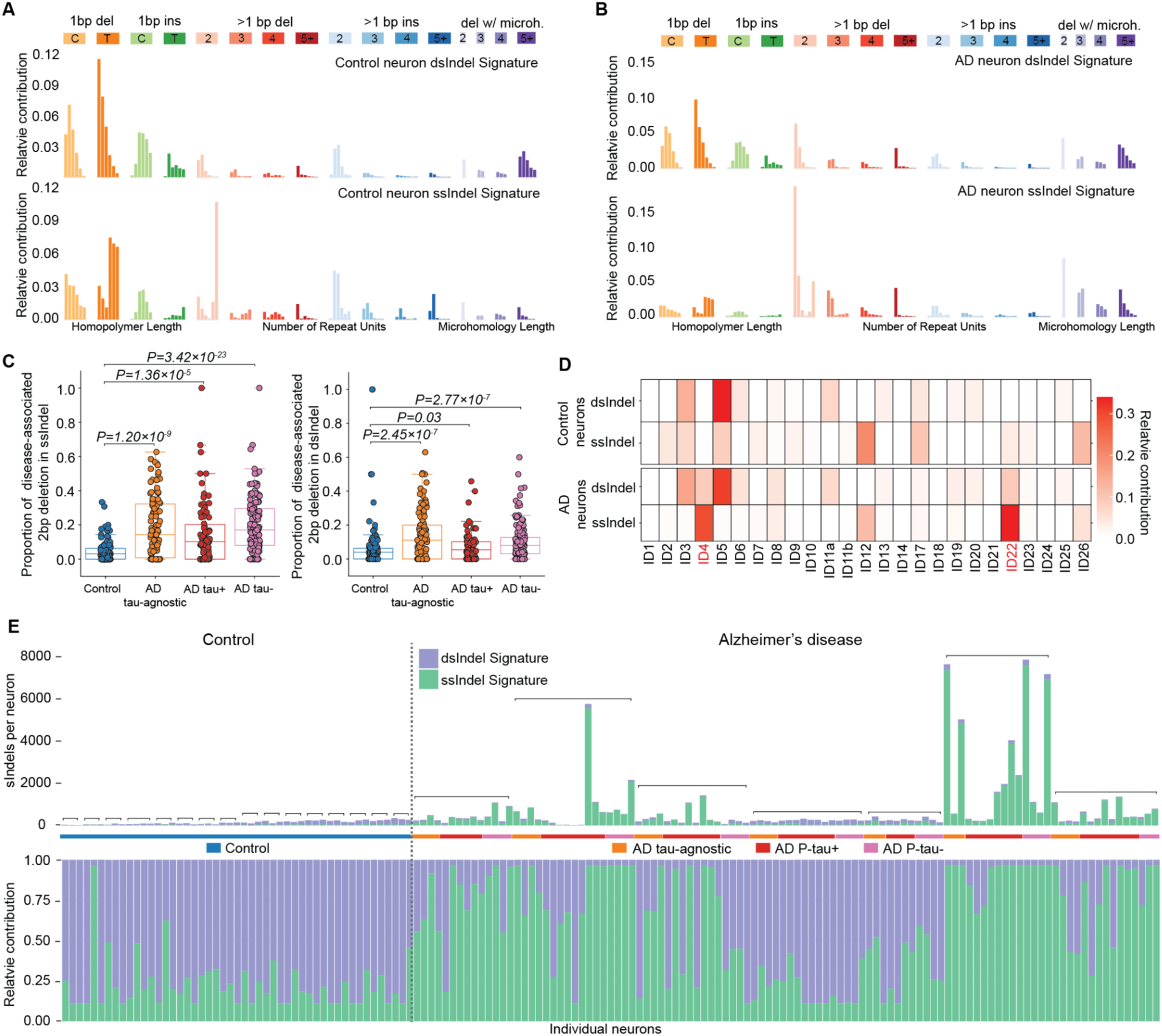
Strand-specific analysis of sIndels in AD neurons. **(A, B)** Strand-resolved sIndel mutational signatures, extracted from META-CS data from control (A) and AD (B) neurons, for double-stranded (dsIndel) and single-stranded (ssIndel) variants. **(C)** Proportion of disease-associated 2bp deletions (occurring at 1-2 repeat units or in microhomologous sequences) in ssIndels and dsIndels from META-CS. **(D)** Decomposition of dsIndel and ssIndel signatures into MuSiCal Indel signatures (ID1-ID26) using NMF. Signatures ID4 and ID22, which contribute to the excessive sIndel burden in AD neurons, contribute to both double- and single-stranded AD Indels. **(E)** Total (top) and relative (bottom) contribution of dsIndel and ssIndel in control and AD neurons. Strand-resolved sIndel signatures extracted from META-CS data were fitted to PTA sIndel spectra to determine the relative contributions of each signature to the overall sIndel burden. Neurons are ordered by increasing age of donor, with neurons from the same individual grouped and denoted with a bracket. For individuals with AD, neurons are organized in the following order: tau-agnostic, tau+, and tau–. A subset of AD neurons shows a pronounced increase in ssIndels.

We then fitted the strand-resolved sIndel signatures to PTA data, enabling us to estimate strand-specific sIndel burdens. Control neuron sIndels were predominantly double-stranded, with consistently low sIndel burden. In contrast, AD neurons varied significantly in sIndel burden, even among neurons from the same individual. Among high-burden AD neurons (genome-wide sIndel burden greater than 300/neuron), single-stranded lesions accounted for on average 88.1% of sIndels. In contrast, sIndels of low-burden AD neurons resembled controls, with the majority being double-stranded (Fig. 5E). Notably, the single-stranded tendency of AD-related variants was specific to sIndels, a distinction from sSNVs.

### NFTs influence AD-related sIndel accumulation through a non-cell autonomous mechanism

Our findings suggested an apparent contradiction: while sIndel accumulation and spectra are independent of single-cell tau pathology, sIndel burden correlates with tissue-wide NFT pathology (Fig. 1C). To reconcile these seemingly divergent results, we hypothesized that mechanisms driving sIndel accrual in AD neurons are influenced by a field effect of upstream AD components that also contribute to quantifiable tissue-wide NFT pathology, which would account for the lack of dependence on single-cell tau state. To test this, we examined how the patterns of sIndels in AD neurons vary as a function of tissue-wide pathologic features of AD (Fig. 6A-B).

**Fig. 6.**
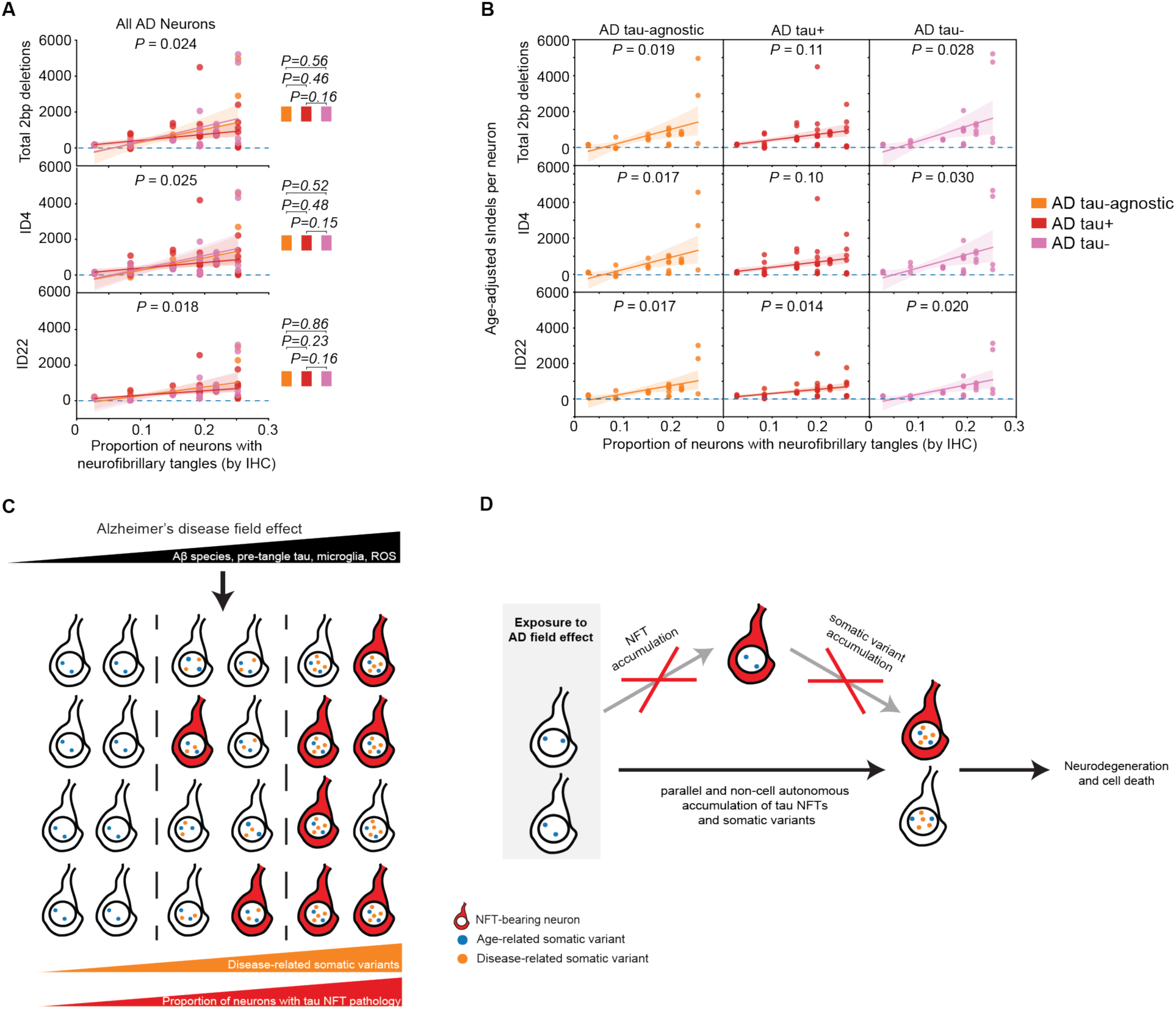
Tissue-wide AD pathology as a contributor to somatic mutation across single-neuron tau states. **(A)** Relationship between tissue proportion of NFT-bearing neurons and sIndel burden by length and ID signature. sIndels are fitted against the proportion of neurons with NFTs, as identified by IHC, using an LME model. The neuronal burden of 2bp deletions and sIndels attributed to ID4 and ID22 is associated with tissue-wide NFT pathology (*P* = 0.024, *P* = 0.025, and *P* = 0.018, respectively). **(B)** Association of tissue-wide NFT pathology with disease-associated sIndel patterns by tau pathologic state. Mechanisms contributing to sIndel accrual are independent of single-cell NFT pathology. **(C)** Proposed non-cell autonomous mechanism of mutagenesis for somatic genomic damage in AD. Non-NFT contributors to AD pathogenesis– including Aβ species, pre-tangle tau, microglia, and ROS–may create an “AD field effect”, driving the parallel yet independent accumulation of both NFTs and somatic mutations. The accumulation of DNA damage may contribute to neurodegeneration and cell death. **(D)** Updated single-cell model of somatic variant and NFT accumulation in AD, suggesting that tau NFTs do not cause somatic genomic changes, but instead that they occur in parallel, non-cell-autonomously, on the pathway to neuron death.

We first observed a positive correlation between the proportion of NFT-bearing neurons and the AD-associated 2bp somatic deletion pattern (*P* = 0.032, Fig. 6A, linear mixed model). We then examined how the mechanisms underpinning 2bp deletions may be influenced by global NFT pathology. We found that the burden of ID4 and ID22 variants–the primary drivers of 2bp somatic deletions–also increase as the proportion of neighboring NFT-bearing neurons rises (Fig. 6A). For all three disease-related sIndel patterns–2bp deletions, ID4, and ID22–we observed similar associations with NFT pathology across AD neurons of all tau states (Fig. 6B). This suggests that the vulnerability to these genotoxic mechanisms is not directly mediated by tau tangle pathology.

## Discussion

Given the robust clinicopathologic role of tau NFTs in AD, we had hypothesized that AD neurons carrying NFT pathology would have elevated somatic mutation burdens. However, using methods to isolate single neurons by their tau NFT pathologic status, we instead found consistently increased somatic mutation burdens in AD neurons across each tested NFT pathologic state. While the substantial accumulation of disease-related somatic mutations in neurons lacking NFT pathology is surprising, this observation aligns with certain findings from a transcriptomic study where differences between NFT-bearing and NFT-free neurons were less pronounced than AD-control differences (*19*).

While NFTs and somatic mutations appear to be decoupled at the single-neuron level, immunohistochemistry analysis revealed that somatic mutations are associated with global (tissue-wide) NFT pathology. This suggests a model whereby various potentially upstream elements of AD, such as amyloid-β, activated microglia, and non-NFT tau species, convey a “field effect” that triggers both tau tangle pathology and somatic mutations, with the latter two events not mechanistically interdependent (Fig. 6C). Future studies to evaluate the effects of upstream constituents of AD pathology on the somatic genome may further illuminate the non-cell-autonomous mechanism of neuronal mutation accrual.

Based on our single-neuron genome findings of somatic mutations across tau NFT pathologic states, we re-examined the sequence of mechanistic events in AD pathogenesis from our previously proposed neuronal model (*12*). The accumulation of disease-related somatic mutations in tau– neurons, on par with tau+ neurons, suggests that tau tangle formation may not precede somatic damage in the genome, but instead that the two processes may occur in parallel in AD (Fig. 6D). Rather than NFTs driving mutagenesis, our new findings suggest a model in which an AD field effect initiates a pathogenic cascade, which includes both the assembly of tau into NFTs and the accrual of genomic damage (Fig. 6D). These somatic genomic changes– particularly 2bp deletions–disrupt genes related to neuronal function and may ultimately contribute to neurodegeneration and cell death.

One implication of our findings is that NFTs appear not to mediate neuronal vulnerability to adverse somatic mutational effects, which would diverge–at least for the genome–from the classical conception of tau as the “executioner” of neurons (*5*). The decoupling of single-cell tau misfolding pathology from somatic mutation accrual accords with observations from a comparative analysis of neuronal subtypes, which suggests a dissociation between NFTs and susceptibility to cell death in AD (*19*). Furthermore, in transgenic mice carrying a MAPT variant encoding pathogenic tau, neurons harboring tangles were less prone to death than neighboring NFT-free neurons, suggesting even a potential protective function for tau tangles (*6*). In this context, we note that we observed marginally lower burdens of sSNVs and sIndels in tau+ than tau– neurons. Together, our findings and these studies suggest that NFTs may not play a significant toxic role in AD. These study discrepancies may potentially suggest the existence of both NFT-dependent and NFT-independent mechanisms of vulnerability in ADs. The pathogenicity of somatic mutations may represent an NFT-independent mechanism that contributes alongside possible NFT effects on neurodegeneration. Considering neurodegenerative conditions beyond AD, these findings may suggest a decoupling of certain intraneuronal protein misfolding pathologies from somatic mutation accumulation, and perhaps more broadly from cellular dysfunction and death.

Pathology (*20*) and imaging (*21*) studies implicate tau as the strongest pathologic staging marker of AD that correlates with cognitive symptoms, consistent with our findings that tissue-level tau NFTs correlate with somatic mutation accrual. However, insofar as genomic somatic mutation provides a window into neuronal health, our findings suggest that tau NFTs may not directly harm neurons in a cell-autonomous fashion. This carries implications for investigational therapeutic strategies targeting tau (*22–24*), raising potential limitations, particularly for approaches targeting NFTs. We should note that non-NFT tau species, such as soluble oligomeric forms, can spread from cell to cell (*25*) and harm neurons (*26*, *27*); (*28*), suggesting that therapeutic strategies that target tau upstream of assembly to full NFTs may be more effective. However, a recent protein structural study suggested that tau oligomers may be composed of the same fibrillar tau as NFTs (*29*), potentially limiting the separate targeting of oligomers and NFTs.

We identified here a specific signature of sIndels, found in a subset of AD neurons but not in controls, primarily consisting of 2bp deletions and characterized by the *de novo* derived mutational signature ID-AD. As with sSNVs, neurons carried these somatic deletions irrespective of tau pathologic states. Our complementary use of strand-agnostic and duplex-strand scWGS methods enabled us to identify that the majority of these somatic deletions are single-stranded, along with a smaller double-stranded component. A subset of these somatic deletions align with the MuSiCal cancer signature ID4, similar to COSMIC ID4, which has been linked to TOP1-mediated ribonucleotide excision repair, suggesting that these AD-related genomic changes may partly result from transcription-associated mutagenesis (*17*). Parallel studies of chronic traumatic encephalopathy (CTE), amyotrophic lateral sclerosis (ALS), and frontotemporal dementia (FTD) have shown similar enrichments in single-stranded 2bp deletions in disease neurons (*30*, *31*). ID-AD thus may be described more aptly as ID-Neurodegeneration (ID-ND) to indicate its presence in a range of neurodegenerative diseases. Moreover, the ALS-FTD study provided experimental support for the potential role of TOP1 in neurodegenerative somatic deletions (*31*). However, in AD, neuronal sIndels of the TOP1-linked ID4 signature showed limited evidence of association with expression, as would be expected for the transcription-associated TOP1 mutagenesis mechanism. Further pointing to mechanisms distinct from TOP1, a substantial proportion of the somatic deletions align with a different signature, MuSiCal ID22. While no specific etiology has been linked to ID22 yet, one report has shown that environmental exposure to nitro-PAH compounds can cause similar somatic deletions (*18*), suggesting a distinct potential mechanism may impact AD pathogenesis. Nonetheless, ID-AD, composed of deletions of at least 2 bases, predominantly at non-repeat sites, appears to represent the most abundant somatic mutation pattern yet identified in neurodegeneration. Although further research is needed to clarify the mechanisms underlying ID-AD, our analysis highlights that the processes contributing to somatic variants are independent of tau deposition at the single-cell level. Moreover, the presence of somatic deletions across tau states and a range of neurodegenerative diseases, even including those associated with misfolded proteins distinct from tau, suggest a more general feature of neurodegeneration and a potential shared therapeutic target.

## Acknowledgments

Human tissue was obtained from the Massachusetts Alzheimer’s Disease Research Center (1P30AG062421-01) and the NIH Neurobiobank at the University of Maryland, and we thank the donors and families for their invaluable contributions to the advancement of science. We thank Ronald Mathieu and Rajesh Krishnan for their assistance with fluorescence-activated nuclear sorting. We thank Jen Neil for input in human subjects. We thank Chris Walsh, Joe Luquette, and Zinan Zhou for helpful discussions, and Andrew Stern for feedback on the manuscript.

## Author Contributions

M.B.M. conceived and designed the study. M.B.M., K.B., S.M.N., S.L.K., and S.P.W. performed single-neuron sorting and single-cell genome sequencing experiments. B.J. performed computational analyses, including PTA and META-CS variant calling as well as burden and signature analyses. K.B. and B.J. optimized experimental and computational parameters for META-CS, with input from M.B.M. and A.Y.H. S.M.N. performed immunofluorescence. B.T.H., D.H.O., M.P.F., and M.B.M. provided clinic-pathologic analysis and selection of disease cases. T.C.S. performed immunohistochemistry. D.S. and E.L.H. analyzed immunohistochemistry and histopathology data, with input from M.B.M. and D.H.O. G.D. contributed to the analysis pipeline and interpretation for duplex sequencing. S.M. contributed to the analysis pipeline for transcriptional and epigenetic enrichment. D.J.S. assisted with data interpretation. M.B.M. and A.Y.H. supervised the study. B.J., K.B., and M.B.M. wrote the manuscript, with input from A.Y.H.

## Data and materials availability

Sequencing data generated in this study will be deposited in the NIH Alzheimer’s disease genomic data public repository, NIAGADS, with controlled use conditions set by human privacy regulations. All of the code used in this study is publicly available in https://github.com/jin-bowen/AD-tau-manuscript. Other materials are available from the authors upon reasonable request.

## Supplementary Materials

Materials and Methods

Figs. S1 to S8

Tables S1 to S6

## References and Notes

1. Alzheimer’s Association. National Institute on Aging-Alzheimer’s Association guidelines for the neuropathologic assessment of Alzheimer’s disease: a practical approach. Acta Neuropathol (2012).

2. R. van der Kant, L. S. B. Goldstein, R. Ossenkoppele, Amyloid-Œ≤-independent regulators of tau pathology in Alzheimer disease. Nat. Rev. Neurosci. 21, 21–35 (2020).

3. S. Dujardin, C. Commins, A. Lathuiliere, P. Beerepoot, A. R. Fernandes, T. V. Kamath, M. B. De Los Santos, N. Klickstein, D. L. Corjuc, B. T. Corjuc, P. M. Dooley, A. Viode, D. H. Oakley, B. D. Moore, K. Mullin, D. Jean-Gilles, R. Clark, K. Atchison, R. Moore, L. B. Chibnik, R. E. Tanzi, M. P. Frosch, A. Serrano-Pozo, F. Elwood, J. A. Steen, M. E. Kennedy, B. T. Hyman, Tau molecular diversity contributes to clinical heterogeneity in Alzheimer’s disease. Nat. Med. 26, 1256–1263 (2020).

4. H. Braak, I. Alafuzoff, T. Arzberger, H. Kretzschmar, K. Del Tredici, Staging of Alzheimer disease-associated neurofibrillary pathology using paraffin sections and immunocytochemistry. Acta Neuropathol. 112, 389–404 (2006).

5. T. Gómez-Isla, R. Hollister, H. West, S. Mui, J. H. Growdon, R. C. Petersen, J. E. Parisi, B. T. Hyman, Neuronal loss correlates with but exceeds neurofibrillary tangles in Alzheimer’s disease. Ann. Neurol. 41, 17–24 (1997).

6. T. J. Zwang, E. D. Sastre, N. Wolf, N. Ruiz-Uribe, B. Woost, Z. Hoglund, Z. Fan, J. Bailey, L. Nfor, L. Buée, K. P. R. Nilsson, B. T. Hyman, R. E. Bennett, Neurofibrillary tangle-bearing neurons have reduced risk of cell death in mice with Alzheimer’s pathology. Cell Rep. 43, 114574 (2024).

7. M. d’Orange, G. Aurégan, D. Cheramy, M. Gaudin-Guérif, S. Lieger, M. Guillermier, L. Stimmer, C. Joséphine, A.-S. Hérard, M.-C. Gaillard, F. Petit, M. C. Kiessling, C. Schmitz, M. Colin, L. Buée, F. Panayi, E. Diguet, E. Brouillet, P. Hantraye, A.-P. Bemelmans, K. Cambon, Potentiating tangle formation reduces acute toxicity of soluble tau species in the rat. Brain 141, 535–549 (2018).

8. L. J. Luquette, M. B. Miller, Z. Zhou, C. L. Bohrson, Y. Zhao, H. Jin, D. Gulhan, J. Ganz, S. Bizzotto, S. Kirkham, T. Hochepied, C. Libert, A. Galor, J. Kim, M. A. Lodato, J. I. Garaycoechea, C. Gawad, J. West, C. A. Walsh, P. J. Park, Single-cell genome sequencing of human neurons identifies somatic point mutation and indel enrichment in regulatory elements. Nat. Genet. 54, 1564–1571 (2022).

9. M. A. Lodato, R. E. Rodin, C. L. Bohrson, M. E. Coulter, A. R. Barton, M. Kwon, M. A. Sherman, C. M. Vitzthum, L. J. Luquette, C. N. Yandava, P. Yang, T. W. Chittenden, N. E. Hatem, S. C. Ryu, M. B. Woodworth, P. J. Park, C. A. Walsh, Aging and neurodegeneration are associated with increased mutations in single human neurons. Science 359, 555–559 (2018).

10. D. Xing, L. Tan, C.-H. Chang, H. Li, X. S. Xie, Accurate SNV detection in single cells by transposon-based whole-genome amplification of complementary strands. Proc. Natl. Acad. Sci. U. S. A. 118, e2013106118 (2021).

11. F. Abascal, L. M. R. Harvey, E. Mitchell, A. R. J. Lawson, S. V. Lensing, P. Ellis, A. J. C. Russell, R. E. Alcantara, A. Baez-Ortega, Y. Wang, E. J. Kwa, H. Lee-Six, A. Cagan, T. H. H. Coorens, M. S. Chapman, S. Olafsson, S. Leonard, D. Jones, H. E. Machado, M. Davies, N. F. Øbro, K. T. Mahubani, K. Allinson, M. Gerstung, K. Saeb-Parsy, D. G. Kent, E. Laurenti, M. R. Stratton, R. Rahbari, P. J. Campbell, R. J. Osborne, I. Martincorena, Somatic mutation landscapes at single-molecule resolution. Nature 593, 405–410 (2021).

12. M. B. Miller, A. Y. Huang, J. Kim, Z. Zhou, S. L. Kirkham, E. A. Maury, J. S. Ziegenfuss, H. C. Reed, J. E. Neil, L. Rento, S. C. Ryu, C. C. Ma, L. J. Luquette, H. M. Ames, D. H. Oakley, M. P. Frosch, B. T. Hyman, M. A. Lodato, E. A. Lee, C. A. Walsh, Somatic genomic changes in single Alzheimer’s disease neurons. Nature 604, 714–722 (2022).

13. P. V. Arriagada, J. H. Growdon, E. T. Hedley-Whyte, B. T. Hyman, Neurofibrillary tangles but not senile plaques parallel duration and severity of Alzheimer’s disease. Neurology 42, 631–639 (1992).

14. V. Gonzalez-Pena, S. Natarajan, Y. Xia, D. Klein, R. Carter, Y. Pang, B. Shaner, K. Annu, D. Putnam, W. Chen, J. Connelly, S. Pruett-Miller, X. Chen, J. Easton, C. Gawad, Accurate genomic variant detection in single cells with primary template-directed amplification. Proc. Natl. Acad. Sci. U. S. A. 118, e2024176118 (2021).

15. ICGC/TCGA Pan-Cancer Analysis of Whole Genomes Consortium, Pan-cancer analysis of whole genomes. Nature 578, 82–93 (2020).

16. H. Jin, D. C. Gulhan, B. Geiger, D. Ben-Isvy, D. Geng, V. Ljungström, P. J. Park, Accurate and sensitive mutational signature analysis with MuSiCal. Nat. Genet. 56, 541–552 (2024).

17. M. A. M. Reijns, D. A. Parry, T. C. Williams, F. Nadeu, R. L. Hindshaw, D. O. Rios Szwed, M. D. Nicholson, P. Carroll, S. Boyle, R. Royo, A. J. Cornish, H. Xiang, K. Ridout, Genomics England Research Consortium, Colorectal Cancer Domain UK 100,000 Genomes Project, A. Schuh, K. Aden, C. Palles, E. Campo, T. Stankovic, M. S. Taylor, A. P. Jackson, Signatures of TOP1 transcription-associated mutagenesis in cancer and germline. Nature 602, 623–631 (2022).

18. J. E. Kucab, X. Zou, S. Morganella, M. Joel, A. S. Nanda, E. Nagy, C. Gomez, A. Degasperi, R. Harris, S. P. Jackson, V. M. Arlt, D. H. Phillips, S. Nik-Zainal, A compendium of mutational signatures of environmental agents. Cell 177, 821–836.e16 (2019).

19. M. Otero-Garcia, S. U. Mahajani, D. Wakhloo, W. Tang, Y.-Q. Xue, S. Morabito, J. Pan, J. Oberhauser, A. E. Madira, T. Shakouri, Y. Deng, T. Allison, Z. He, W. E. Lowry, R. Kawaguchi, V. Swarup, I. Cobos, Molecular signatures underlying neurofibrillary tangle susceptibility in Alzheimer’s disease. Neuron 110, 2929–2948.e8 (2022).

20. B. T. Hyman, C. H. Phelps, T. G. Beach, E. H. Bigio, N. J. Cairns, M. C. Carrillo, D. W. Dickson, C. Duyckaerts, M. P. Frosch, E. Masliah, S. S. Mirra, P. T. Nelson, J. A. Schneider, D. R. Thal, B. Thies, J. Q. Trojanowski, H. V. Vinters, T. J. Montine, National Institute on Aging-Alzheimer’s Association guidelines for the neuropathologic assessment of Alzheimer’s disease. Alzheimers. Dement. 8, 1–13 (2012).

21. R. Ossenkoppele, G. Salvadó, S. Janelidze, A. Pichet Binette, D. Bali, L. Karlsson, S. Palmqvist, N. Mattsson-Carlgren, E. Stomrud, J. Therriault, N. Rahmouni, P. Rosa-Neto, E. M. Coomans, E. van de Giessen, W. M. van der Flier, C. E. Teunissen, E. M. Jonaitis, S. C. Johnson, S. Villeneuve, PREVENT-AD Research Group, T. L. S. Benzinger, S. E. Schindler, R. J. Bateman, J. D. Doecke, V. Doré, A. Feizpour, C. L. Masters, C. Rowe, H. J. Wiste, R. C. Petersen, C. R. Jack Jr, O. Hansson, Plasma p-tau217 and tau-PET predict future cognitive decline among cognitively unimpaired individuals: implications for clinical trials. Nat. Aging, doi: 10.1038/s43587-025-00835-z (2025).

22. A. Mullard, Anti-tau antibody stumbles in phase II Alzheimer trial. Nat. Rev. Drug Discov. 23, 883 (2024).

23. C. J. Mummery, A. Börjesson-Hanson, D. J. Blackburn, E. G. B. Vijverberg, P. P. De Deyn, S. Ducharme, M. Jonsson, A. Schneider, J. O. Rinne, A. C. Ludolph, R. Bodenschatz, H. Kordasiewicz, E. E. Swayze, B. Fitzsimmons, L. Mignon, K. M. Moore, C. Yun, T. Baumann, D. Li, D. A. Norris, R. Crean, D. L. Graham, E. Huang, E. Ratti, C. F. Bennett, C. Junge, R. M. Lane, Tau-targeting antisense oligonucleotide MAPTRx in mild Alzheimer’s disease: a phase 1b, randomized, placebo-controlled trial. Nat. Med. 29, 1437–1447 (2023).

24. E. E. Congdon, C. Ji, A. M. Tetlow, Y. Jiang, E. M. Sigurdsson, Tau-targeting therapies for Alzheimer disease: current status and future directions. Nat. Rev. Neurol. 19, 715– 736 (2023).

25. B. Frost, R. L. Jacks, M. I. Diamond, Propagation of tau misfolding from the outside to the inside of a cell. J. Biol. Chem. 284, 12845–12852 (2009).

26. B. Hyman, All the tau we cannot see. Annu. Rev. Med. 74, 503–514 (2023).

27. C. A. Lasagna-Reeves, D. L. Castillo-Carranza, U. Sengupta, A. L. Clos, G. R. Jackson, R. Kayed, Tau oligomers impair memory and induce synaptic and mitochondrial dysfunction in wild-type mice. Mol. Neurodegener. 6, 39 (2011).

28. G. Niewiadomska, W. Niewiadomski, M. Steczkowska, A. Gasiorowska, Tau oligomers neurotoxicity. Life (Basel) 11, 28 (2021).

29. S. Lövestam, D. Li, J. L. Wagstaff, A. Kotecha, D. Kimanius, S. H. McLaughlin, A. G. Murzin, S. M. V. Freund, M. Goedert, S. H. W. Scheres, Disease-specific tau filaments assemble via polymorphic intermediates. Nature 625, 119–125 (2024).

30. G. Dong, C. C. Ma, S. Mao, S. M. Naik, K. S.-M. Brown, G. A. McDonough, J. Kim, S. L. Kirkham, J. D. Cherry, M. Uretsky, E. Spurlock, A. C. McKee, A. Y. Huang, M. B. Miller, E. A. Lee, C. A. Walsh, Diverse somatic genomic alterations in single neurons in chronic traumatic encephalopathy, bioRxiv.org (2025). 10.1101/2025.03.03.641217.

31. Z. Zhou, L. J. Luquette, G. Dong, J. Kim, J. Ku, K. Kim, M. Bae, D. D. Shao, B. Sahile, M. B. Miller, A. Y. Huang, W. J. Nathan, A. Nussenzweig, P. J. Park, C. Lagier-Tourenne, E. A. Lee, C. A. Walsh, Recurrent patterns of widespread neuronal genomic damage shared by major neurodegenerative disorders, bioRxiv.org (2025). 10.1101/2025.03.03.641186.

32. P. Bankhead, M. B. Loughrey, J. A. Fernández, Y. Dombrowski, D. G. McArt, P. D. Dunne, S. McQuaid, R. T. Gray, L. J. Murray, H. G. Coleman, J. A. James, M. Salto-Tellez, P. W. Hamilton, QuPath: Open source software for digital pathology image analysis. Sci. Rep. 7, 16878 (2017).

33. U. Schmidt, M. Weigert, C. Broaddus, G. Myers, Cell detection with star-convex polygons, arXiv [cs.CV] (2018). http://arxiv.org/abs/1806.03535.

34. C. L. Bohrson, A. R. Barton, M. A. Lodato, R. E. Rodin, L. J. Luquette, V. V. Viswanadham, D. C. Gulhan, I. Cortés-Ciriano, M. A. Sherman, M. Kwon, M. E. Coulter, A. Galor, C. A. Walsh, P. J. Park, Linked-read analysis identifies mutations in single-cell DNA-sequencing data. Nat. Genet. 51, 749–754 (2019).

35. F. Manders, A. M. Brandsma, J. de Kanter, M. Verheul, R. Oka, M. J. van Roosmalen, B. van der Roest, A. van Hoeck, E. Cuppen, R. van Boxtel, MutationalPatterns: the one stop shop for the analysis of mutational processes. BMC Genomics 23, 134 (2022).

36. M. Petljak, L. B. Alexandrov, J. S. Brammeld, S. Price, D. C. Wedge, S. Grossmann, K. J. Dawson, Y. S. Ju, F. Iorio, J. M. C. Tubio, C. C. Koh, I. Georgakopoulos-Soares, B. Rodríguez-Martín, B. Otlu, S. O’Meara, A. P. Butler, A. Menzies, S. G. Bhosle, K. Raine, D. R. Jones, J. W. Teague, K. Beal, C. Latimer, L. O’Neill, J. Zamora, E. Anderson, N. Patel, M. Maddison, B. L. Ng, J. Graham, M. J. Garnett, U. McDermott, S. Nik-Zainal, P. J. Campbell, M. R. Stratton, Characterizing mutational signatures in human cancer cell lines reveals episodic APOBEC Mutagenesis. Cell 176, 1282– 1294.e20 (2019).

37. L. B. Alexandrov, J. Kim, N. J. Haradhvala, M. N. Huang, A. W. Tian Ng, Y. Wu, A. Boot, K. R. Covington, D. A. Gordenin, E. N. Bergstrom, S. M. A. Islam, N. Lopez-Bigas, L. J. Klimczak, J. R. McPherson, S. Morganella, R. Sabarinathan, D. A. Wheeler, V. Mustonen, PCAWG Mutational Signatures Working Group, G. Getz, S. G. Rozen, M. R. Stratton, PCAWG Consortium, The repertoire of mutational signatures in human cancer. Nature 578, 94–101 (2020).

38. S. M. A. Islam, M. Díaz-Gay, Y. Wu, M. Barnes, R. Vangara, E. N. Bergstrom, Y. He, M. Vella, J. Wang, J. W. Teague, P. Clapham, S. Moody, S. Senkin, Y. R. Li, L. Riva, T. Zhang, A. J. Gruber, C. D. Steele, B. Otlu, A. Khandekar, A. Abbasi, L. Humphreys, N. Syulyukina, S. W. Brady, B. S. Alexandrov, N. Pillay, J. Zhang, D. J. Adams, I. Martincorena, D. C. Wedge, M. T. Landi, P. Brennan, M. R. Stratton, S. G. Rozen, L. B. Alexandrov, Uncovering novel mutational signatures by de novo extraction with SigProfilerExtractor. Cell Genom. 2, None (2022).

39. K. Wang, M. Li, H. Hakonarson, ANNOVAR: functional annotation of genetic variants from high-throughput sequencing data. Nucleic Acids Res. 38, e164 (2010).

