## Supplementary material for "Neurons accumulate disease-specific somatic genomic changes across tau pathologic states in Alzheimer’s disease": Materials and Methods, Supplementary Figures S1-S8

**The PDF file includes:**

Materials and Methods
Supplementary Text
Figs. S1 to S8
Tables S1 to S6

### Materials and Methods

Human tissue samples and selection of cases

Post-mortem frozen human tissues were obtained from the Massachusetts Alzheimer's Disease Research Center (MADRC) at Massachusetts General Hospital and the NIH Neurobiobank at the University of Maryland Brain and Tissue Bank (UMBTB). Tissue collection and distribution for research and publication was conducted according to protocols approved by the MassGeneral Brigham (MGB) Institutional Review Board (for MADRC: 1999P009556) and the University of Maryland Institutional Review Board (for UMBTB: 00042077), after provision of written authorization and informed consent. Research on these de-identified specimens and data was performed at Brigham and Women's Hospital with MGB Institutional Review Board approval (secondary-use non-human subjects protocol 2019P003790). A set of neurotypical control and Alzheimer's disease neuron data analyzed here were generated as part of a previous study (*12*). Neurotypical control cases had no clinical history of dementia or other neurological disease, and the age-matched control subset was defined as age > 50 years. AD cases were ascertained for a clinical history of dementia consistent with AD, pathologically confirmed high-stage AD neuropathologic change (Braak stage V–VI), and no other significant neurodegenerative pathologies.

Isolation of pyramidal neurons for single-cell studies by FANS

Single neuronal nuclei were isolated from frozen (-80 °C) unfixed postmortem human dorsolateral prefrontal cortex (dlPFC) brain tissue, using FANS for the neuronal nuclear transcription factor (NeuN), as described in previous work (*12*) and shown in Fig. 1D.

For PTA experiments, nuclei were isolated beginning with homogenization of brain tissue in a dounce homogenizer using a chilled tissue lysis buffer (10 mM Tris-HCl, 0.32 M sucrose, 3 mM Mg(OAc)2, 5 mM CaCl2, 0.1 mM EDTA, 1 mM DTT, 0.1% Triton X-100, pH 8) on ice. Tissue homogenates were layered on top of a sucrose cushion buffer (1.8 M sucrose 3 mM Mg(OAc)2, 10 mM Tris-HCl, 1 mM DTT, pH 8) and ultra-centrifuged for 1 h. at 30,000 × g. Nuclear pellets were resuspended in ice-cold PBS supplemented with 3 mM $MgCl_2$, filtered, then stained.

For META-CS experiments, nuclei were isolated from brain tissue in a dounce homogenizer using a chilled tissue lysis buffer (0.25 M sucrose, 5 mM $MgCl_2$, 25 mM KCl, 10 mM Tris-HCl, pH 8, 0.1% Triton X-100, 1 µM DTT) on ice. Tissue homogenates were filtered through a 40 µm cell strainer and centrifuged for 10 min. at 900 × g at 4 °C. Nuclear pellets were resuspended in ice-cold immunostaining buffer (PBS, 1% BSA) and centrifuged for 5 min. at 500 × g. Pellets were resuspended in ice-cold immunostaining buffer, filtered, and stained.

For all scWGS experiments, nuclear suspensions were stained with DAPI and anti-NeuN (RBFOX3) antibody directly conjugated to Alexa Fluor 488 (AF488) (Millipore MAB377X, clone A60, 1:1250). DAPI staining allowed for the isolation of single diploid neuronal nuclei apart from debris and doublet droplets. FANS was used to distinguish and sort single neuronal nuclei by their NeuN immunofluorescence signal. FANS was performed using a BD FACS Aria II (software BD FACSDiva version 8.0.2/8.0.3) and analyzed with FlowJo (version 10.4). PBS was used as sheath fluid with a sheath pressure of 20 psi and a 100-µm nozzle.

NeuN staining produced a bimodal signal distribution. Large neuronal nuclei, representing excitatory pyramidal neurons, were then identified by targeting the nuclei with highest NeuN signal among the NeuN+ neuronal fraction, while also gating for the population with the highest forward scatter area (FSC-A) signal. As we reported previously, this high-FSC-A, high-NeuN population represents large pyramidal neurons, which are >99% excitatory neurons (*12*), comprising 2–5% of the total population of nuclei in each sample.

Separation of tau pathologic state by FANS

To study specific populations by their tau pathologic state, during FANS, pyramidal neurons (isolated as described above) were distinguished and further sorted by their tau NFT pathology immunofluorescence signal, into tau+ and tau– nuclei (shown in Fig. 1D). Three anti-tau antibodies were used, in separate experiments, each directly conjugated to AF647 for visualization. The majority of experiments utilized phospho-specific anti-P-tau(Ser404) (2Z4G) antibody conjugated to Alexa Fluor 647 (AF647) (Cell Signaling Technology 98523, 1:1000), while a subset used phospho-specific anti-P-tau(T205) (E7D3E) antibody conjugated to Alexa Fluor 647 (AF647) (Cell Signaling Technology, 53001, 1:250) and total anti-tau (HT7) antibody conjugated to AF647 (Invitrogen, 51-5916-42, 1:200) for comparison. Anti-tau antibody signal showed a cluster of tau– nuclei with a scattering of high-staining nuclei sorted as tau+ (Fig. 1D). Tau+ populations from the three antibodies showed comparable findings (fig. S2); therefore analyses focused on the combined group of tau+ neurons. Tau– nuclei refer to those sorted negative for anti-tau(Ser404)-AF647. A mixture analysis determined the false-positive rate as <0.3%. To distinguish from neurons sorted for tau+ or tau– status, we refer to AD neurons not profiled for tau pathology as "tau-agnostic" AD neurons.

Single-cell genome sequencing of neurons using PTA

Single neurons, prepared as described above, were sorted one-nucleus-per-well into 96-well plates and their genomes were amplified using PTA (*8*), a method that pairs an isothermal DNA polymerase with a termination base to achieve quasi-linear amplification. PTA reactions were performed using the ResolveDNA Whole Genome Amplification Kit v1 (BioSkryb Genomics). Nuclei were sorted into 3 µL Cell Buffer pre-chilled on ice. Nuclei were then lysed by addition of 3 µL MS Mix, with mixing at 1,400 rpm performed after each step. Notably, this lysis step was performed on ice (distinct from some protocols that perform lysis at room temperature). Lysed nuclei were then neutralized with 3 µL SN1 buffer, followed by 3 µL of SDX reagent and a 10-min. incubation at room temperature. Next, 8 µL of reaction mix (containing polymerase) was then added, for a total reaction volume of 20 µL. Amplification was carried out for 10 h. at 30 °C, followed by enzyme inactivation at 65 °C for 3 min. Amplified DNA was then cleaned up using AMPure XP beads (Beckman Coulter; A63880), and the yield was determined using PicoGreen binding (Quant-iT dsDNA Assay Kit, Thermo Fisher Scientific). Of note, experiments were performed with reagents not containing EvaGreen dye, which was available in some kit versions to track product yield fluorescently but carries risk of somatic mutation artifacts. Samples were then subjected to quality control by multiplex PCR for four random genomic loci as previously described (*9*). Amplified genomes showing positive amplification for all four multiplex PCR loci were prepared for Illumina sequencing.

Libraries were prepared following a modified KAPA HyperPlus Library Preparation protocol described in the ResolveDNA EA Whole Genome Amplification protocol. In brief, end repair and A-tailing were performed for 100-500 ng amplified DNA input. Adapter ligation was then performed using the SeqCap Adapter Kit (Roche, 07141548001). Ligated DNA was cleaned up using AMPure and amplified through an on-bead PCR amplification. Amplified libraries were selected for a size of 300–600 bp using AMPure. Libraries were subjected to quality control using PicoGreen and TapeStation HS DS100 Screen Tape (Agilent PN 5067-5584) before sequencing. Single-cell genome libraries were sequenced on the Illumina NovaSeq platform (150 bp × 2) at 30× coverage (Table S1).

Duplex single-cell genome sequencing of neurons using modified META-CS

The genomes of single neurons were amplified by a method based on META-CS (*10*), a transposase-based whole-genome amplification technique in which each DNA fragment is tagged and barcoded with 16 unique tags (Dataset S1 of (*10*)), allowing for single-cell, strand-resolved identification. DNA oligonucleotides were obtained from Integrated DNA Technologies (IDT) with HPLC purification. Each of the 16 META-CS DNA transposons were annealed and assembled into transposomes with Tn5 transposase (Diagenode Tagmentase, C01070010) per manufacturer's protocol and stored at -80 °C.

Single neuronal nuclei, isolated as previously described, were sorted one-per-well into 96-well plates, and lysed in 2 µL of 1x single-cell lysis buffer (20 mM Tris, pH 8.0, 20 mM NaCl, 0.15% Triton X-100, 25 mM dithiothreitol, 1 mM EDTA, 15 mg/mL thermolabile proteinase K (TLPK) (NEB, P8111S)) at 30 °C for 1 h., then 55 °C for 10 min. for protease inactivation. The TLPK concentration used in the original protocol was increased tenfold to improve yield and library diversity upon amplification (Fig. S7). Single-cell lysates were stored at -20 °C if not immediately processed for amplification.

Lysed nuclei were then transposed by the addition of 8 µL transposition mix (5ul Diagenode 2X Tagmentation buffer (Diagenode; C01019043), 1 µL diluted META-CS transposome pool, 2 µL $H_2O$), mixed at 1640 rpm for 1 min., spun down at 1500 rpm, and incubated at 55 °C for 15 min. Transposases were removed by the addition of 2 µL 6X stop buffer containing 300 mM NaCl, 45 mM EDTA, 0.01% Triton X-100, and 1 mg/mL TLPK, with mixing and incubation at 37 °C for 30 min., followed by protease inactivation at 55 °C for 10 min.

First-strand tagging was performed by the addition of 13 µL strand-tagging mix 1 containing 5 µL Q5 reaction buffer (NEB; B9027S), 5 µL Q5 high GC enhancer (NEB; B9028A), 0.85 µL 100 µM (total) Adp1 primer mix (Dataset S1 of (*10*)), 0.6 µL 100 mM MgCl2, 0.55 µL water, 0.5 µL 10 mM (each) dNTP mix (Thermo Scientific; R0192), 0.25 µL of 20 mg/mL bovine serum albumin (NEB; B9000S), and 0.25 µL Q5 DNA polymerase (NEB; M0491S), followed with mixing and incubation at 72 °C for 3 min., 98 °C for 30 s., 62 °C for 5 min., 72 °C for 1 min. Adp1 primers were removed with 1 µL thermolabile exonuclease I (NEB; M0568L), with mixing and incubation at 37 °C for 15 min., followed by exonuclease inactivation at 65 °C for 5 min.

Second-strand tagging was performed by the addition of 4 µL strand-tagging mix 2 containing 1 µL Q5 reaction buffer, 1 µL Q5 high GC enhancer, 0.95 µL 100 µM (total) Adp2 primer mix (Dataset S1 of (*10*)), 0.85 µL water, 0.1 µL 10 mM (each) dNTP mix, and 0.1 µL Q5 DNA polymerase, and incubation at 72 °C for 3 min., 98 °C for 30 s., 62 °C for 5 min., 72 °C for 1 min. Adp2 primers were removed by repeating the exonuclease step described above.

Strand tagging products were amplified by the addition of 19 µL PCR mix containing 2 µL NEBNext 96 Index Primers (NEB; E6609S), 4 µL Q5 reaction buffer, 4 µL Q5 high GC enhancer, 0.4 µL 10 mM (each) dNTP mix, 8.4 µL water, and 0.2 µL Q5 DNA polymerase and incubation at 98 °C for 20 s., 11-13 cycles of [98 °C for 10 s., 72 °C for 2 min.], 72 °C for 2 min.

Single-cell libraries were pooled together, and then libraries were purified by DNA Clean and Concentrator-5 columns (Zymo; D4013) and amplification efficiency was checked for fragment size and concentration by Agilent TapeStation. Size selection was performed with Ampure, wherein the pooled library was divided into three groups based on fragment size. Medium-size fragments (~300 - 1000 bp) were selected first by the addition of 0.5X beads then by a further addition of 0.25X beads (for a final 0.75X).

Single-cell libraries were sequenced on the Illumina NovaSeqX platform (150 bp × 2), targeting at least 30× coverage per 10-cell pool (Table S1).

Quantifying NFT-bearing neurons by IHC

Five-micron formalin-fixed paraffin-embedded (FFPE) brain sections from the superior frontal gyrus were immunostained for phospho-tau (epitope Ser202, Thr205) using a monoclonal antibody (AT8) (ThermoFisher, Waltham, MA, Cat# MN1020, 1:5000) and for Aβ (DAKO, Santa Clara, CA. 6F/3D, 1:600) on a Bond Rx autostainer (Leica Biosystems, Buffalo Grove, IL), with one parallel staining run for tau AT8 IHC used for quantification of samples in this study. 3,3'-diaminobenzidine (DAB) brown chromogen was used for signal detection. Slides were digitally scanned (Hamamatsu NanoZoomer) at 40X setting (400X total magnification). Digital images were analyzed using QuPath (version 0.5.1) (*32*). Three representative regions of cortex per slide were manually annotated, and 2D cell detection was performed using the StarDist plugin (*33*).. A random forest model was trained across the dataset on cells manually annotated to specifically identify large pyramidal neurons (10 per slide). The model was applied to all images along with a DAB-channel threshold to quantify both DAB-negative and DAB-positive neurons (i.e., those bearing tau neurofibrillary tangles).

Detection of somatic SNVs and Indels from PTA data with SCAN2

We first used bwa-mem2 (version 2.2.1) to align reads from bulk and PTA fastq files to the human genome GRCh37(hg19) (with decoy of hs37d5). We then used MarkDuplicates and base quality score recalibration (bsqr) from the Genome Analysis Toolkit (version 4.1.9.0) to mark duplicate reads and rescale base pair quality.

We used a modified version of SCAN2 (*8*) to identify sSNVs and sIndels from PTA data. The modified SCAN2, available at https://github.com/jin-bowen/SCAN2, demonstrates comparable performance to the original version, while reducing computing time by utilizing intermediate files for joint calling and cross-sample construction. When tested on control neurons, the mutation rates for sSNVs and sIndels show high correlations between the modified and original SCAN2 (0.989 for sSNV and 0.955 for sIndel). Similar results were found in AD tau-agnostic neurons too (0.987 for sSNV and 0.997 for sIndel).

The modified SCAN2 uses the command “scan2 config” and “scan2 run” with individual PTA bam files and matched Bulk WGS bam files to call sSNVs and sIndels. Of note, we used the gatk3_gvcf mode to enable joint calling with intermediate gVCF files that could be used to generate a Panel of Normals later. For “scan2 config”, we used common variants from dbSNP (version 20180423) to filter out potentially contaminated germline variants, and the 1000 Genomes Project Phase 3 phasing panel for haplotype phasing.

The cross-sample panel that filtered false positives in sIndels was generated with intermediate gVCF files. We used the command "joint_panel" to create the Panel of Normals for control, AD tau-agnostic, tau+, and tau– neurons, respectively. Subsequently, we updated the joint calling results with the cross-sample panel to flag false-positive sIndels.

To ensure the accuracy of the mutational burden estimated by SCAN2, we cross-referenced it with an independent sSNV caller, LiRA (*34*). LiRA employs an independent method to identify sSNVs, based on read-level haplotype phasing with nearby germline heterozygous polymorphisms. This approach leverages the linkage of reads to phase somatic mutations accurately. One neuron was excluded due to the low concordance of the estimated sSNV burden between LiRA and SCAN2. Neurons passing QC and included in this study are included in Table 1.

Mixed-effects modeling of somatic mutation burden

We employed a linear mixed-effects model with *mixedlm* from *statsmodels* (version 0.14.0) to estimate the age-burden association. We used the following formula: *burden ~ age + group + (1/case)*, where the group is the type of neuron, including control, AD tau-agnostic, tau+, and tau–. Case refers to the case ID for neurons from the same individual. As such, intra-individual variability is incorporated in the mixed effect and would not drive the association test results.

Excess somatic mutation burden was calculated by subtracting the normal aging-related somatic mutation accumulation observed in control neurons. To establish the age-burden relationship for control neurons, we used a mixed-effect model with *mixedlm* from *statsmodels* (version 0.14.0). The slope (control) and intercept (control) derived from this model represent normal aging patterns in neurotypical individuals.

Subsequently, the mutation burdens of AD neurons—including tau-agnostic, tau+, and tau– neurons—were calculated using the following formula: *excess burden = burden - age * slope(control) - intercept(control)*. This approach isolates the excess burden in AD neurons from age-related somatic mutation burden, to reflect disease-specific factors.

Mutational signature analysis

1) Fitting sSNV spectra with known signatures: First, sSNVs were classified into 96 SBS types, to generate a spectrum representing all types of neurons, including control, AD tau-agnostic, tau+, and tau–. We used the *MutationPatterns* package (*35*) (version 3.8.1) function *fit_to_signatures* to generate the linear combination of given mutational signatures that most closely reconstruct the original spectra. We inputted the function with Signatures A,C, SBS scE and SBS scF. Signatures A and C were previously identified from single-cell WGS of human neurons (*9*). Signature A has been found in non-disease neurons and associated with aging, Signature C is disease-associated and contains excess C>A substitutions (*12*). SBS scE and SBS scF are artifact signatures reported to be associated with multiple displacement amplification (MDA) (*36*), and we observed negligible contribution in PTA-amplified neuronal genomes in this study.

2) *De novo* signature analysis for sIndels: First, sIndels were classified into 83 types (ID83 classification) (*37*), enabling the generation of spectra that encompass all the types of neurons, including control, AD tau-agnostic, AD tau+, and AD tau–. We used non-negative matrix factorization (NMF) to decompose the spectra. To determine the optimal rank, we tested values

ranging from 1 to 10. The rank that maximized the cophenetic correlation coefficient and showed a clear inflection point in the residual sum of squares curve was 2. As a result, we decomposed the spectrum into two *de novo* signatures. The de novo signature analysis was performed using *SigProfilerExtractor (38)* (version 1.2.28).

3) Signature decomposition for sIndels: We used SigProfilerAssignment to assign custom Indel signatures to the given spectra, inputting COSMIC signatures into the tool. However, *de novo* signature ID-AD did not match any single COSMIC signature or their linear combinations, based on a cosine similarity threshold of 0.8. To explore further, we also inputted MuSiCal signatures, which represent a refined set of signatures identified by the MuSiCal algorithm (*16*). MuSiCal signatures ID1–14, 17, and 18 show high similarity (cosine similarity > 80%) to COSMIC ID signatures, and MuSiCal includes 9 distinct Indel signatures without direct analogues in COSMIC.

Mutational enrichment expression analysis

Mutational enrichment analysis was conducted by permuting somatic variants and examining their distribution across genic regions. For total mutational enrichment, all variants identified from PTA data were randomly shuffled across the genome using the SCAN2 permute command. This permutation was repeated 10,000 times to generate 10,000 null sets of mutations, while mutation types remained fixed during the process. For genic regions, enrichment was assessed as the ratio of observed mutations (directly called mutations) to expected mutations (randomly permuted mutations). For each gene, the average expression value was derived from published snRNA-seq data in excitatory neurons from prefrontal cortex (*30*). Based on these expression levels, genes were categorized into 10 deciles, with decile 1 representing the least expression and decile 10 representing the greatest expression. We tested the relationship between gene expression decile and mutation enrichment using a linear model.

Signature-specific mutational enrichment analyses followed a similar approach, with two differences. First, after permutation, each null set of mutations was decomposed into predefined mutational signatures. When quantifying enrichment, we compared the observed mutations, which are decomposed into specific signatures, to the expected mutations, which were permuted and also decomposed into the same signatures.

Functional impact of somatic Indels

We use ANNOVAR (*39*) (version 20150322) to predict sIndel consequence and to estimate genome-wide high-impact sIndels. First, we used ANNOVAR to annotate all observed sIndels. We classified sIndels that caused frameshifts as high-impact Indels. We extrapolated the genome-wide high-impact Indel burden by the sensitivity of SCAN2 given for each neuron.

Identification of sSNVs and sIndels from META-CS data

1) Preprocessing of META-CS data: Data for somatic SNV and Indel calling was processed by a previously described computational pipeline (*30*), which extends the original META-CS pipeline that was designed exclusively for somatic SNV calling (*10*). First, we used the pre-meta module to preprocess paired-end reads, which involved trimming Illumina adapters, extracting Tn5 barcodes, and merging overlapping read ends. Next, we mapped the reads to the human reference

genome (hs37d5) using both BWA-MEM (version 0.7.17) and minimap2 (version 2.26-r1175). Neurons were excluded if they showed excessively uneven coverage (coefficient of variation, CoV, greater than 10). Cells passing QC are included in Table 1. The BAM files for individual neurons were further split by unique pairs of Tn5 barcodes, as reads with the same pairs of Tn5 barcodes originate from the same DNA fragments. The order of Tn5 barcodes, determined by the primer at the strand tagging stage, carries strand-specific information from the original DNA fragments.

2) Detection of double-stranded SNVs (dsSNVs) and Indels (dsIndels): To ensure robust and accurate variant calling, we applied several detection and filtering criteria. For dsSNVs, we excluded reads with more than three mismatches. For candidate dsSNVs and dsIndels, we filtered out sites present among gnomAD SNPs and Indels that have a population frequency of 1% or greater. For a candidate to be considered a true mutation, it must meet the following criteria: no alternative allele reads present in the bulk reference genome sample, a minimum of four total reads mapped to the candidate locus (a4), and at least two alternative reads from each strand (s2) to provide duplex-strand support in the cell. Additionally, the variant allele frequency (VAF) must be above 0.2 for the mutation to be considered true (0.9 for SNV), considering reads from both strands. Neurons were excluded if they showed insufficient coverage, as indicated by absence of any mutation calls.

3) Detection of single-stranded SNVs (ssSNVs) and Indels (ssIndels): For ssSNVs, we similarly excluded reads with more than 3 mismatches. ssSNV and ssIndel candidates were filtered out if present among gnomAD SNPs and Indels that have a population frequency of 1% or greater. To be considered a true ssSNV or ssIndel, the variant must meet the following criteria: no alternative allele reads present in the bulk reference genome, at least four alternative reads from one strand (s4), and no alternative reads and at least four reference reads from the other strand (s4).

Analysis of somatic variant burdens with double-stranded and single-stranded signatures

First, the spectra for double-strand and single-strand sIndels in control and AD neurons were determined from META-CS data, respectively. Since these spectra were not orthogonal to each other, they could not be used directly for refitting. To address this, we used NMF to decompose the double-strand and single-strand sIndel spectra into distinct double-stranded and single-stranded sIndel signatures with *MutationalPatterns (35)* (version 3.8.1). As a result, these two intermediate signatures became independent of each other and were suitable for refitting. To estimate the proportions of double-strand and single-strand mutations within the PTA call sets, each mutation type in each cell was fitted to the double-stranded and single-stranded sIndel signatures. To estimate the contribution of known signatures to the double-stranded and single-stranded sIndel signatures, we decomposed these signatures into COSMIC signatures using the *fit_to_signatures* function from *MutationalPatterns* (version 3.8.1).

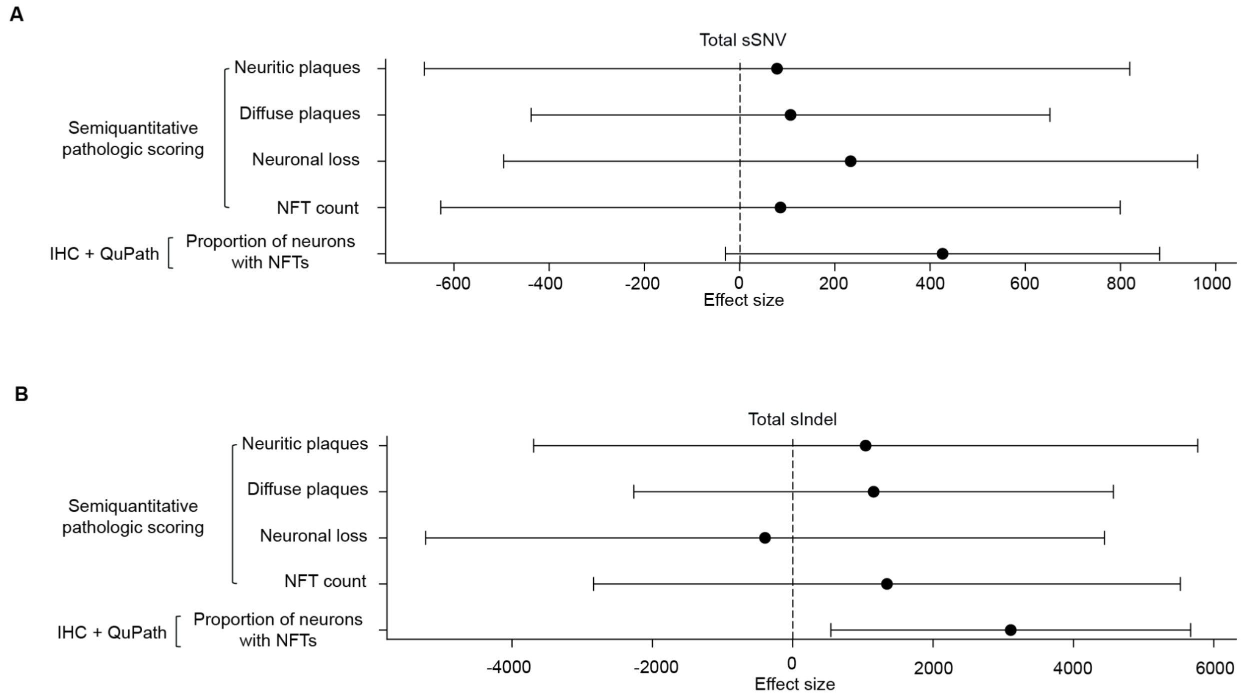


**Fig. S1. Relationship between AD pathologic features and neuronal somatic mutation burden.** sSNV **(A)** and sIndel **(B)** mutation classes were profiled. The association between pathologic features of AD pathology and neuronal somatic variant burdens was evaluated by a mixed linear model. Four semi-quantitative pathologic scoring features were included in the model: diffuse amyloid-β plaques, neuritic amyloid-β plaques, neuronal loss, and tau neurofibrillary tangles (NFTs), each extracted from standardized Alzheimer's disease research center (ADRC) neuropathology reports. Tau NFTs were further profiled in individual cells, enabling a quantitative assessment of the proportion of NFT-bearing neurons (using immunohistochemistry for anti-phospho-tau antibody AT8 and QuPath). All pathologic variables were normalized to better compare the effect size (arbitrary units) across features. The mean and 95% confidence interval are shown.

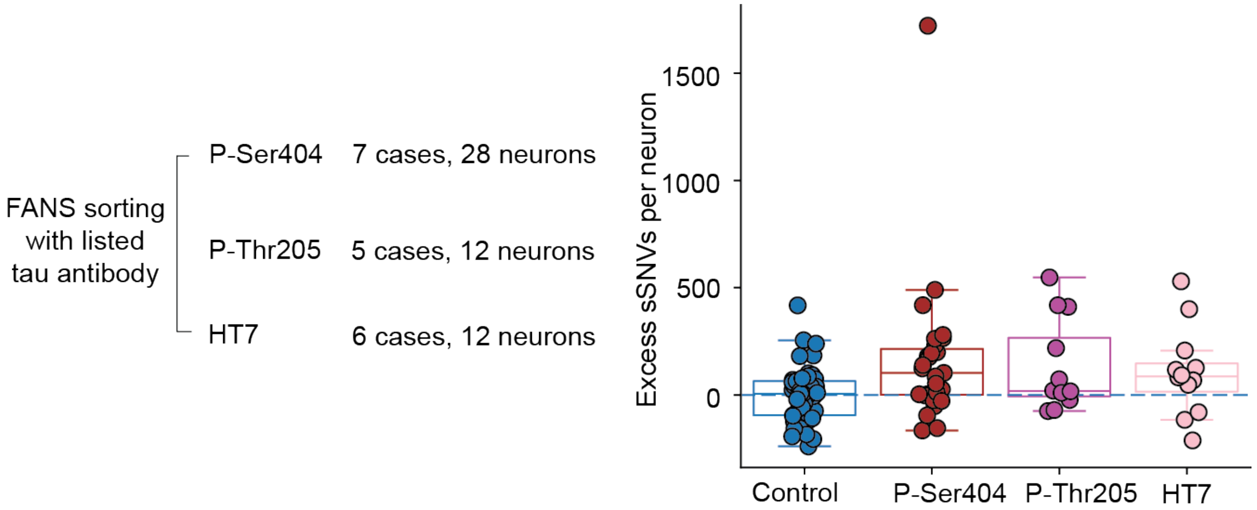


**Fig. S2. Somatic SNV and Indel burdens in tau+ neurons isolated by different tau antibodies.** Three distinct anti-tau antibodies were used to isolate NFT-bearing neurons by FANS: phosphoepitope-targeting antibodies P-Ser404 and P-Thr205 and total tau antibody HT7. For all 7 AD cases, 4 neurons per case were profiled with P-Ser404. Two P-Thr205 tau+ neurons were profiled from 6 selected AD cases. Two HT7 tau+ neurons were profiled from 5 selected AD cases. Excess somatic variants per neuron (sSNV and sIndel separately) were calculated as the residual value from linear regression for age (as in Fig. 2B). Tau+ neurons did not significantly differ in sSNV or sIndel burden between antibodies; each tau+ neuron group showed evidence of increased somatic variants relative to control neurons.

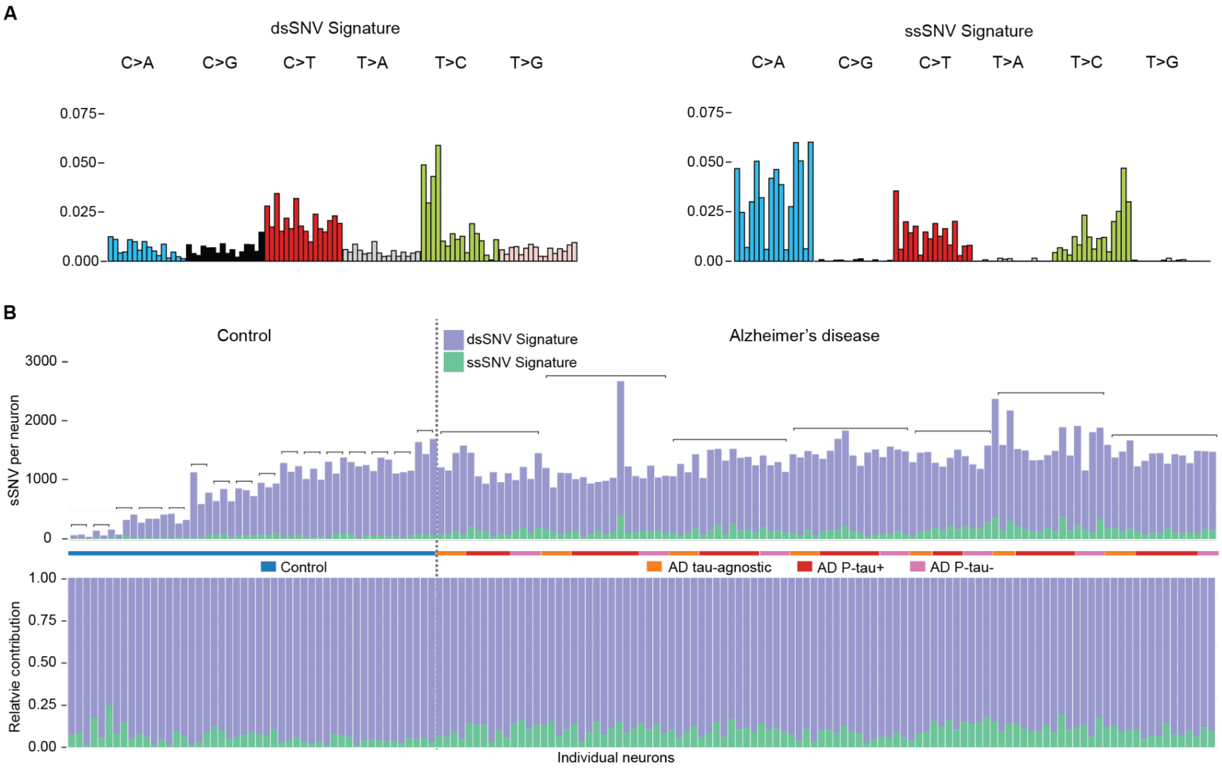


**Fig. S3. Duplex-strand scWGS analysis of somatic SNVs in neurons.** Duplex-strand scWGS (META-CS) was performed on neurons from control, AD tau+, AD tau–, and AD tau-agnostic samples, then called sSNVs for double-stranded mutations (dsSNVs) and single-stranded lesions (ssSNVs). **(A)** Mutational signature analysis produced dsSNV and ssSNV signatures. **(B)** Absolute (upper panel) and relative (lower panel) contributions of dsSNV and ssSNV signatures to PTA scWGS sSNV mutation burdens in control and AD neurons. Individuals are arranged by age, and neurons from the same donor are grouped and marked with a bracket. For AD donors, neurons are organized in the following order: tau-agnostic, tau+, and tau–. Overall, PTA-identified SNVs in neurons are predominantly derived from the double-stranded signature (estimated dsSNV contribution by group: control: 93.7%, AD tau-agnostic: 90.7%, AD tau+: 89.8%,  AD tau–: 88.5%, overall: 91.0%).

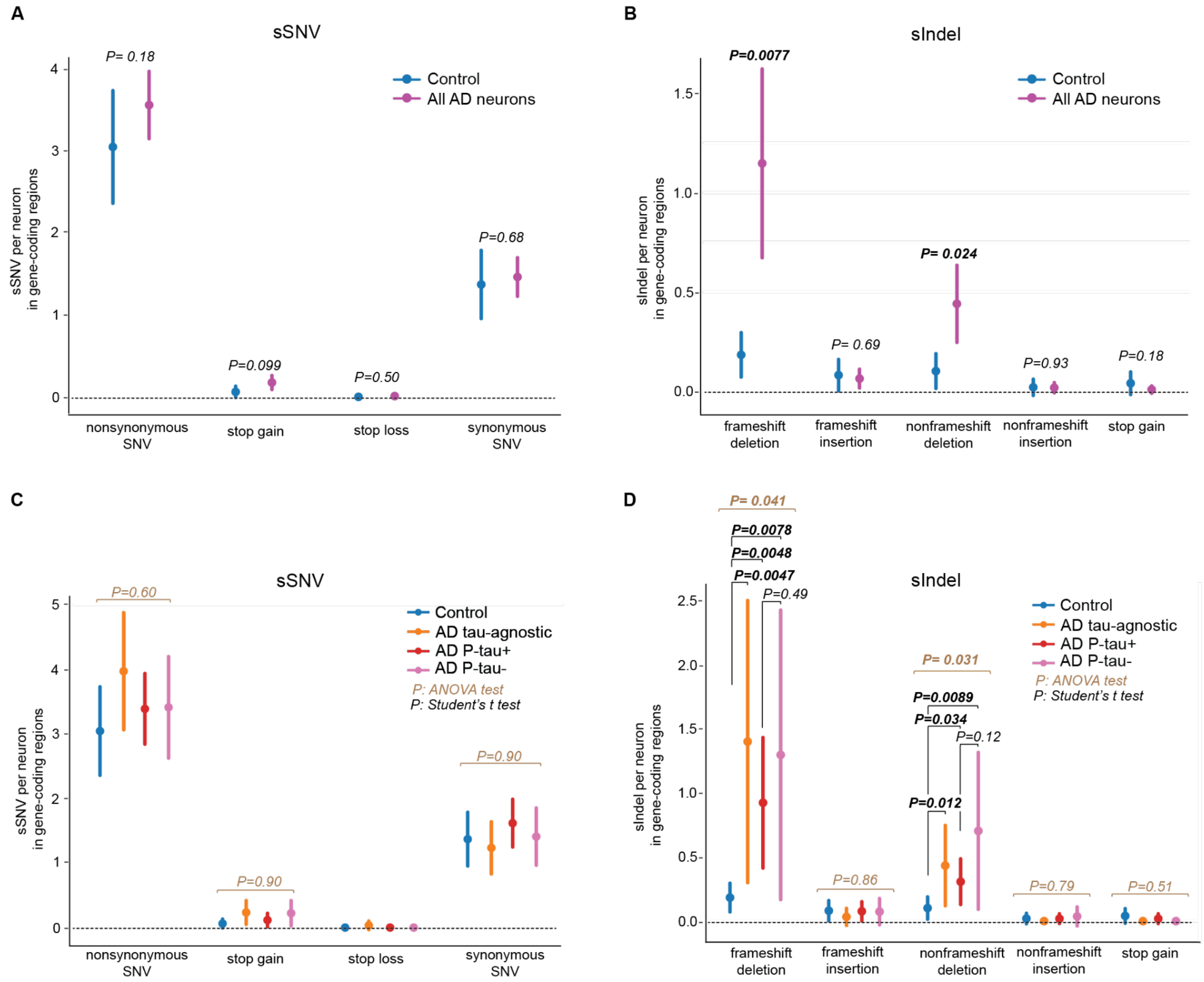


**Fig. S4. Functional analysis of somatic SNVs and Indels in control and AD neurons. (A, B)** Comparison between control and AD neurons, which represent combined data from AD tau+, tau–, and tau-agnostic neurons. The burdens of impactful sSNVs (A) and sIndels (B) within gene-coding regions were compared between AD and control neurons with the Student's t-test. **(C, D)** Comparison across all neuron groups. The burdens of impactful sSNVs (C) and sIndels (D) in gene-coding regions were evaluated and compared among all groups of neurons with ANOVA test (P-values shown when not significant). For groups with a significant ANOVA result, pair-wise Student's t-test comparisons were performed between each group of AD neurons and control neurons, and between tau+ and tau–. All populations of AD neurons, including tau+, tau–, and tau-agnostic, exhibit a higher number of frameshift and non-frameshift deletions relative to control neurons. No significant differences are observed between AD tau+ and tau– neurons.

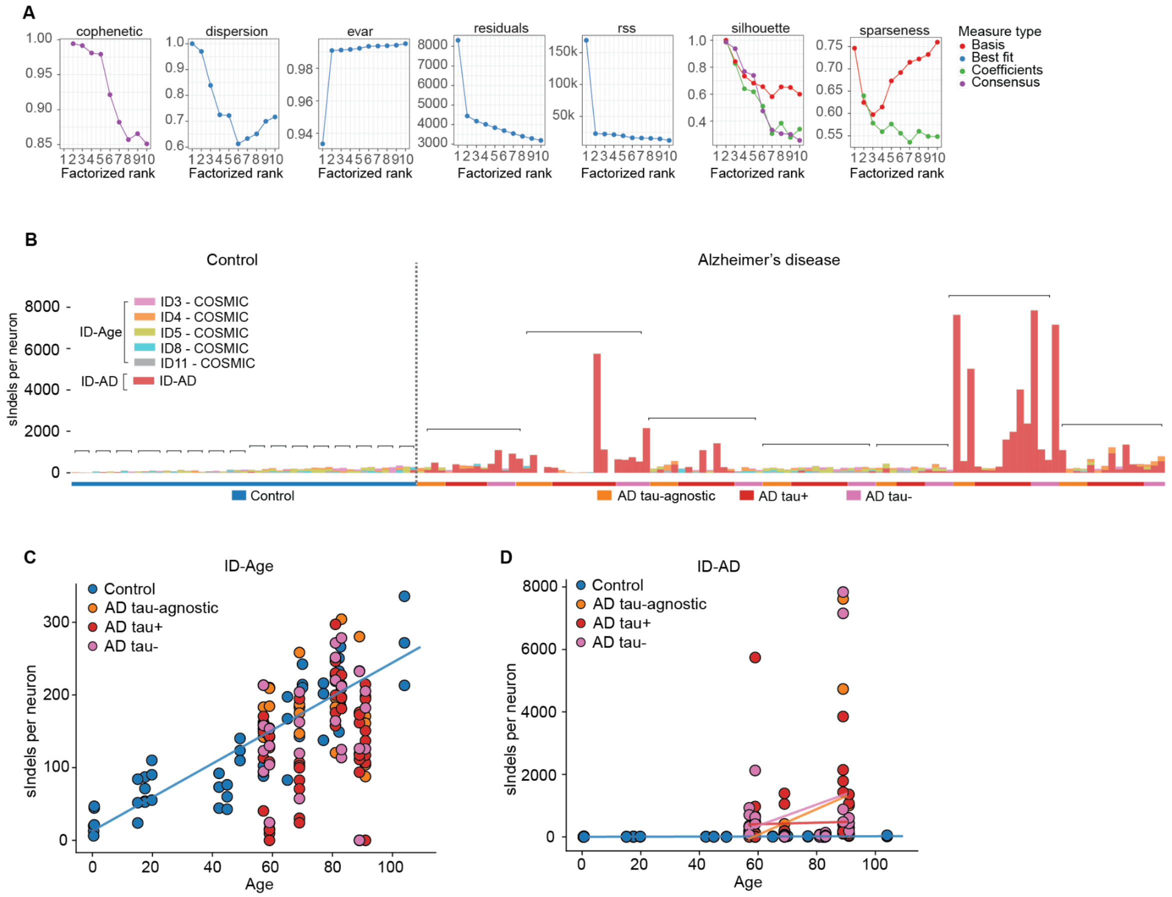


**Fig. S5. *De novo* mutational signature analysis of somatic Indels in control and AD neurons. (A)** Metrics for *de novo* signature extraction. An ID83-style sIndel classification spectra derived from PTA scWGS data was decomposed with NMF analysis with various numbers of the rank (1-10). The smallest rank that maximizes the cophenetic correlation coefficient and exhibits obvious inflection in the residual sum of squares curve is 2. Two signatures, designated ID-AD and ID-Age, were extracted from the PTA spectra. **(B)** Decomposition of neuronal sIndel signatures with COSMIC cancer signatures resolves ID-Age into distinct components and shows novel signature ID-AD as AD-related. Brackets identify neurons from each individual, with colors denoting the tau pathologic state for AD neurons. **(C-D)** sIndel *de novo* mutational signature analysis, showing contributions of ID-Age and ID-AD to sIndel burden by age.

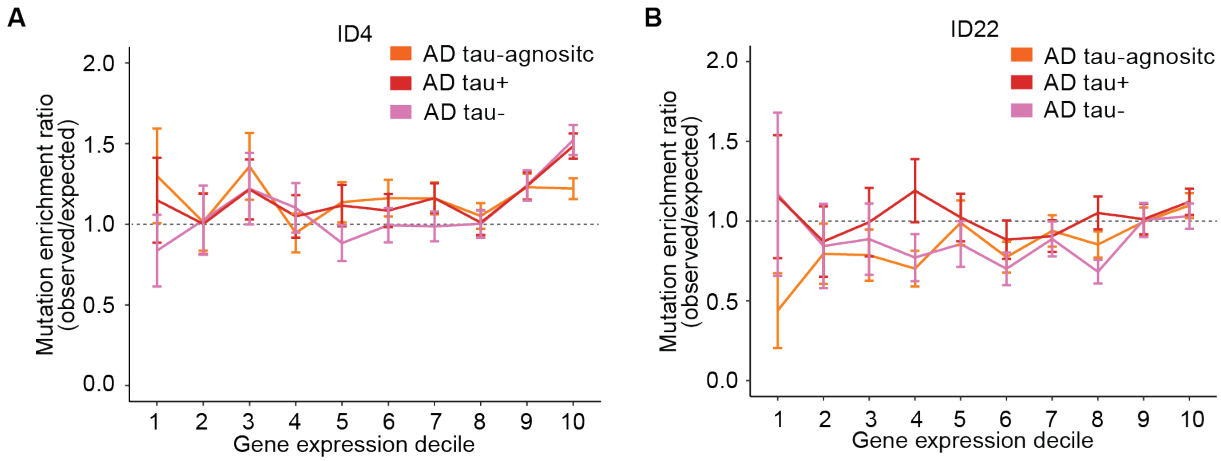


**Fig. S6. Influence of transcriptional activity on somatic Indel accrual in AD neurons, by specific populations. (A-B)** Genomic sIndel density as a function of gene expression, ascertained by scRNAseq from prefrontal cortex excitatory neurons. Indel density is distinguished by MuSiCal mutational signatures ID4 (A) and ID22 (B).

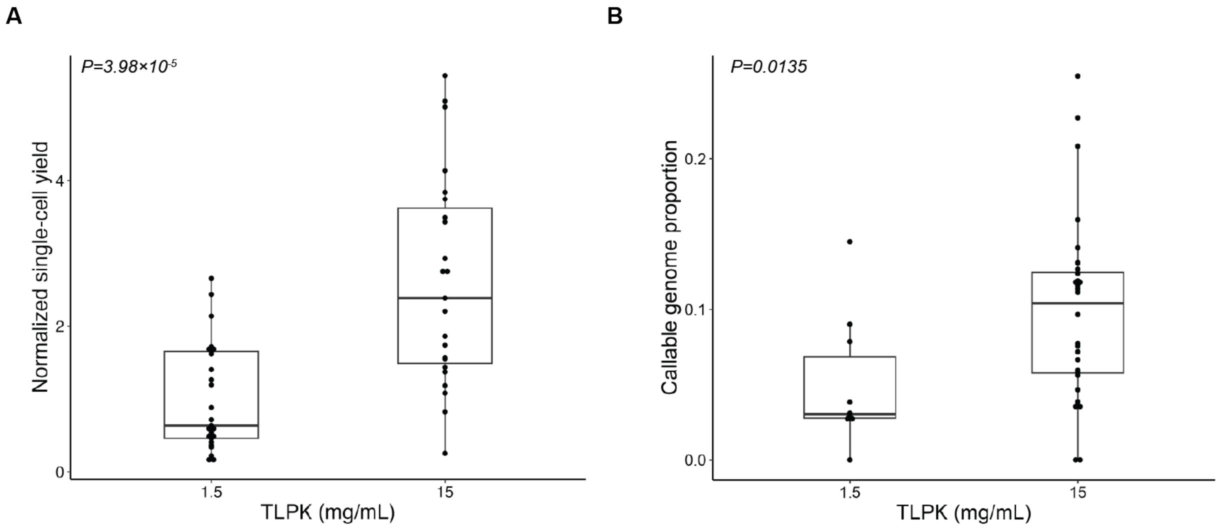


**Fig. S7. Increased protease concentration during META-CS lysis improves single-cell amplified genomic DNA yield and library diversity. (A)** Single-cell META-CS yields following lysis in the original (1.5 mg/mL thermolabile proteinase K, TLPK) and modified (15 mg/mL TLPK) lysis buffer. A tenfold increase in TLPK improves yields and was used for all META-CS data (*P* = $3.9810^{-5}$, two-tailed Wilcoxon test). Single-cell yields are normalized to the mean yield obtained using the original lysis buffer. **(B)** Proportion of the genome per single cell that meets a calling strand threshold of four reads, with at least two reads on each strand (a4s2). A tenfold increase in TLPK significantly increases a4s2 coverage and is indicative of improved library diversity (*P* = 0.0135, two-tailed Wilcoxon test).

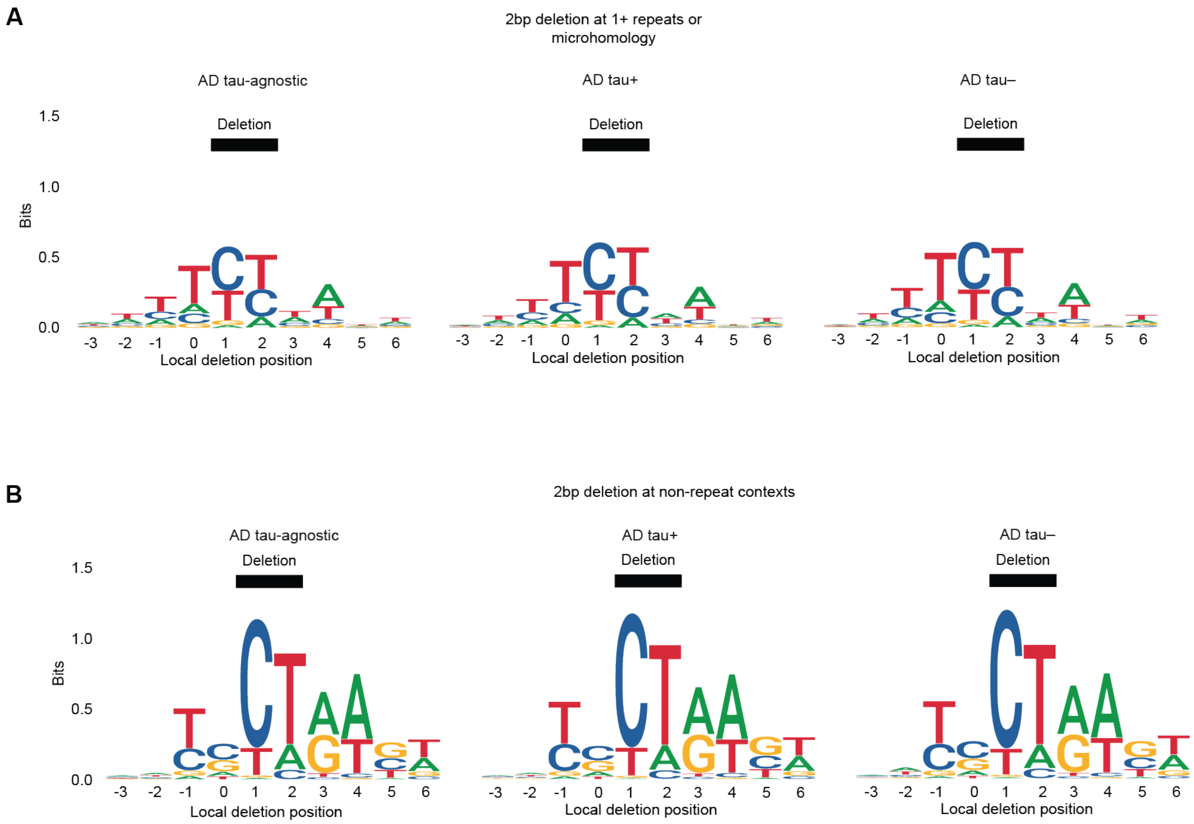


**Fig. S8. Motif analysis of somatic 2bp deletions in repeat- and non-repeat genomic contexts in AD tau-agnostic, tau+, and tau– neurons. (A)** Motif analysis of 2bp deletions in regions with at least one repeat (at least 2 repeat "units") or with microhomology in AD neurons with distinct tau pathologic states. 2bp deletions in repeat regions are concentrated at TNT independently of NFT pathology. **(B)** Motif analysis of 2bp deletions in regions with no repeats (1 repeat "unit") in AD neurons with distinct tau pathologic states. 2bp deletions in non-repeat regions exhibit an NT motif regardless of NFT pathology.

**Table S1. (separate file)**
Sample information. Clinical data, library and sequencing metrics.

**Table S2. (separate file)**
PTA and META-CS sequencing statistics.

**Table S3. (separate file)**
PTA sSNV and sIndel rates.

**Table S4. (separate file)**
PTA sSNV and sIndel calls.

**Table S5. (separate file)**
META-CS sSNV and sIndel calls.

**Table S6. (separate file)**
Gene Ontology terms enriched for sSNVs and sIndels.
